# Single-subject EEG measurement of interhemispheric transfer-time for the in-vivo estimation of axonal morphology

**DOI:** 10.1101/2022.12.19.521040

**Authors:** Rita Oliveira, Marzia De Lucia, Antoine Lutti

## Abstract

Assessing axonal morphology in-vivo opens new avenues for the combined study of brain structure and function. A novel approach has recently been introduced to estimate the morphology of axonal fibers from the combination of MRI data and EEG measures of the interhemispheric transfer time (IHTT). In the original study, the IHTT measures were computed from EEG data averaged across a group, leading to bias of the axonal morphology estimates.

Here, we seek to estimate axonal morphology from individual measures of IHTT, obtained from EEG data acquired in a visual evoked potential experiment. Subject-specific IHTTs are computed in a data-driven framework with minimal *a priori* constraints, based on the maximal peak of neural responses to visual stimuli within periods of statistically significant evoked activity in the inverse solution space. The subject-specific IHTT estimates ranged from 8 to 29 ms except for one participant and the between-session variability was comparable to the differences in IHTT between subjects. The scale parameter of the axonal radius distribution, computed from the IHTT estimates and the MRI data, ranged from 0 to 0.79 *μ*m. The change in axonal g-ratio with axonal radius ranged from 0.62 to 0.81 *μm*^−*α*^.

The single-subject measurement of the IHTT yields estimates of axonal morphology that are consistent with histological values. However, improvement of the repeatability of the IHTT estimates is required to improve the specificity of the single-subject axonal morphology estimates.

## 1 Introduction

The radius and myelin thickness of axons are morphological features that play an essential role in neuronal communications and consequently, brain function (Rushton, 1951; Waxman and Bennett, 1972; MacKay and Laule, 2016). Axons of different radius are affected differentially over the course of several diseases such as multiple sclerosis (Evangelou et al., 2001), autism (Wegiel et al., 2018), and motor neuron disease (Cluskey and Ramsden, 2001). Therefore, the specific assessment of morphological properties of axons from in-vivo data is essential for the study of brain function, and for the monitoring of disease evolution in patient populations. Biomarkers computed from MRI data can allow the in-vivo monitoring of microscopic properties of brain tissue (Lutti et al., 2014; Stikov et al., 2015; Does, 2018). However, these biomarkers suffer from a lack of specificity with regards to morphological properties of axons, and represent averages across the large axon populations present within an MRI voxel.

Recently, we proposed a new approach that allows the non-invasive estimation of axonal radius and relative myelination of axonal fibers in humans (Oliveira et al., 2022). The proposed method relies on a biophysical model that links these morphological properties of white matter fibers and data collected in-vivo. This in-vivo data consists of MRI measures of the g-ratio (gMRI, Campbell et al., 2018) – sampled along a white matter tract of interest – and a measure of axonal conduction velocity in the same tract. In Oliveira et al., 2022 we focused on the visual transcallosal tract, whose conduction velocity can be estimated with an electroencephalography (EEG) paradigm that measures the delay of visual information transfer across hemispheres, *i*.*e*. the Interhemispheric Transfer Time (IHTT).

The bedrock of visual IHTT estimation is the crossed organization of the visual system: a unilaterally-presented visual stimulus first reaches the contralateral cortex and is then transferred to the ipsilateral cortex (Saron and Davidson, 1989; Chaumillon et al., 2018) via the splenium of the corpus callosum (*e*.*g*. Saron and Davidson, 1989; Martin et al., 2007). Due to its temporal resolution, EEG is particularly suited for non-invasive estimation of the IHTT. Classically, this estimation is based on the latency difference of the evoked activity recorded from homologous electrodes (*e*.*g*. Saron and Davidson, 1989; Brown et al., 1994). This approach relies on the assumption that maxima in the voltage spatial distribution across the electrode montage reflect the maxima of the underlying electromagnetic field produced by active neurons. An *ad hoc* selection of the electrodes of interest and of the latency at which the transfer is most likely to occur is required for this approach. Either the positive (P1) or negative (N1) components of the event-related potentials (ERPs) have been considered as a proxy for the neuronal activations (*e*.*g*. Saron and Davidson, 1989; Brown et al., 1994; Ipata et al., 1997; Westerhausen et al., 2006; Moes et al., 2007; Whitford et al., 2011). The P1 component represents an early stage of visual processing (Martin et al., 2007; Whitford et al., 2011), while N1 may be a closer reflection of callosal transfer (Brown and Jeeves, 1993; Ipata et al., 1997). Moreover, while P1 originates from spatially localized activations over the extrastriate cortex of the fusiform gyrus (Ipata et al., 1997; Di Russo et al., 2001), the voltage generators for N1 are widespread, encompassing occipital and parietal regions (Ipata et al., 1997; Di Russo et al., 2001). In the absence of sufficient knowledge of the spatio-temporal distributions of neural activations in response to visual stimuli, the *ad hoc* selection of electrodes and latencies remains an open question.

In this light, our previous work aimed to improve IHTT estimation by proposing an alternative source-based IHTT estimation approach that overcomes the *a priori* choice of electrodes and components (Oliveira et al., 2022). Performing source reconstruction on the voltage measurements enhances EEG spatial resolution. It also allows the reconstruction of neuronal activity within homologous cortical regions and consequently, the computation of the IHTT from the latency difference between the two maximal neuronal activations. In Oliveira et al., 2022 however, estimation of the IHTT was conducted from a group average of the time courses of the source reconstructed signal. According to simulation results, this may have led to bias of up to 40% in the resulting subject-specific estimates of axonal radius and myelination (Oliveira et al., 2022). Thus, estimation of the IHTT at the subject level is of the utmost importance for the accurate estimation of axonal morphological features with this approach.

In the current study, we seek to obtain subject-specific estimates of the IHTT from EEG data to allow accurate estimation of axonal morphology at the subject level. We extend the cortical region considered for the estimation of the maximal activity upon visual stimulation. This enables the inclusion of activity originating from widespread generators of the visual response and mitigates the effect of errors in the localization of the signal sources that have more impact in subject-level analyses. We introduce three quantitative metrics that help resolve ambiguities on the selection of the maximal activity in each hemisphere and support the attribution of the IHTT estimates to visual-evoked activity. We evaluate the within- and between-session repeatability of the IHTT estimates. Finally, we use the subject-specific IHTT values to compute estimates of axonal morphology in the occipital transcallosal tract of each participant. The inter- and intra-subject variabilities of the morphological estimates were compared to assess the specificity of axonal morphological estimates.

## 2 Methods

### 2.1 Participants

The same dataset as in Oliveira et al., 2022 was used. In brief, the data includes a set of EEG and MRI recordings obtained from 17 healthy volunteers. Three participants were excluded due to artifacted EEG recordings, resulting in a final dataset of 14 participants (8 males; age: mean±s.d.=27.14±3.86 years). All participants were right-handed as assessed with the Edinburgh Handedness Inventory (Oldfield, 1971) and three of them were left-eye dominant as evaluated by looking through a small opening at a distant object (Miles, 1930). The participants did not have neurological/psychiatric disorders and had normal or corrected-to-normal vision. Three of these participants were invited to do a second EEG recording session for a test-retest evaluation.

### 2.2 Electroencephalography-Based Estimation of the Interhemispheric Transfer Time

#### 2.2.1 Experimental paradigm

The experimental paradigm is the same as previously reported by our group (Oliveira et al., 2022). We presented a black and white checkerboard on a grey background presented on the lower left visual field (LVF) or right visual field (RVF). The checkboard (4° diameter) was presented on a grey background (20 cd/m^2^) for 100 ms at 6° horizontal and 6° vertical distance from the centrally-located fixation cross (0.8° in size). Each experimental block contained 205 trials (95 for each visual field and 15 with no stimulus). Inter-trial intervals varied randomly from one to two seconds and the trials were presented in a pseudorandom order. The experiment was administered using Psychtoolbox-3.0.16 in Matlab (R2019b, The Mathworks, Natick, MA). Participants were seated 80 cm away from a screen (61 cm, 60 Hz refreshing rate). Subjects were instructed to keep their gaze on the central fixation cross and press a button (with index finger) as quickly as possible after each stimulus appearance. There were 6 experimental blocks presented pseudo-randomly, three were answered with the right hand and the remaining three with the left hand.

#### 2.2.2 Data acquisition

Continuous 128-channel EEG was recorded with an Ag/AgCl electrode cap (waveguard™original, ANT Neuro, Hengelo, Netherlands) and the Micromed recording system (Micromed SystemPlus Evolution, Mogliano Veneto, Italy). All electrodes were referenced to FPz and grounded at AFFz. Electrooculogram (EOG) was recorded using two additional horizontal electrodes placed to the outer canthi of each eye. The sampling rate was 1024 Hz. Electrode impedance was below 20 kΩ. Using the xensor™ digitizer (ANT Neuro, Hengelo, Netherlands) the 3D head shape and electrode positions were digitized for each participant.

#### 2.2.3 Data processing

To preprocess the data, we used Fieldtrip, version 20191206 (Oostenveld et al., 2011) and EEGLAB, version 13.4.4b (Delorme and Makeig, 2004) toolboxes on MATLAB (R2021a, The Mathworks, Natick, MA). We first bandpass-filtered (digital filters) the continuous raw EEG signal at 0.1-40 Hz and extracted EEG epochs from 100 ms prior to stimulus onset to 300 ms after stimulus presentation. Trials containing large eye movements were first removed on visual inspection. Next, we used Independent Component Analysis with the *runica* algorithm (Bell and Sejnowski, 1995) to identify and remove components containing eye movement-related artifacts. Semi-automatic artifact detection with a threshold of 80 *μ*V was used to identify and exclude additional artifact epochs from further analysis: 20.5% and 21.3% of the trials were rejected on average across participants, leading to an average number of accepted trials of 453 (range: 350 to 528) and 449 (range: 364 to 528) for the LVF and RVF stimulation, respectively. The same threshold of 80 *μ*V was used to identify artifact electrodes, which were next interpolated using the nearest neighbor: 5.9% of electrodes (range 3 to 14 electrodes) were interpolated on average across participants. The EEG epochs were re-referenced to the average reference and DC drift was removed by subtracting the average amplitude within each epoch. The global field power (GFP), a measure of the spatial standard deviation of the electrical potentials, was computed for each dataset in order to relate the evoked components with the interhemispheric transfer time latencies (Hamburger and Michelle, 1991).

For source reconstruction, we used Brainstorm, version 16-04-2021 (Tadel et al., 2011) toolbox. Each subject’s MRI-based head was first registered to the EEG-digitized head with an iterative algorithm. A head model using a 3-layer Boundary Element Method (Kybic et al., 2005; Gramfort et al., 2010) was created on each subject MRI image with 15000 vertices on the grey matter. Finally, current densities (CD, pA.m) time courses were estimated for each condition (LVF; RVF), source vertice, and trial for each subject using the minimum-norm current density approach (Hämäläinen and Ilmoniemi, 1994). We defined the occipital region of interest (ROI) as including the *inferior parietal, lateral occipital, superior parietal, cuneus, lingual, fusiform, pericalcarine, precuneus* of the Desikan-Killiany atlas (Desikan et al., 2006). The use of a bigger cortical region at the subject-specific level compared to the primary and secondary visual cortices used originally (Oliveira et al., 2022) is motivated by several reasons. First, the localization of neuronal generators in deep cortical fissures and the specificities of each individual’s cortex make the source reconstruction around the calcarine sulcus challenging (Creel, 2012). Consequently, mislocalization of the signal between hemispheres or across neighboring regions within the same hemisphere is likely to occur (Cuffin, 1998; Grech et al., 2008), with a stronger impact on IHTT estimates at the subject-than group-level (lower signal-to-noise ratio). Second, the widespread distribution of generators over the posterior cortex is consistent with the late latencies of maximal cortical activity used for IHTT estimation (>140 ms). In particular, between 136 and 146 ms (late P1 component), the neuronal generators are localized in the ventral extrastriate cortex of the fusiform gyrus, and from 150 to 200 ms (N1 complex), deep sources can be found in the parietal lobe (Di Russo et al., 2001). Therefore, given that the visual response is not restricted within the primary visual areas, by adopting a larger region we include the activity originating from the main generators of such response.

In order to ensure that neuronal activity was due to the evoked activity in response to the visual stimuli, we identified the post-stimulus period during which the estimated CDs were statistically different compared to the baseline period (Supp. Fig. 1a). To this aim, we implemented a cluster permutation statistical test (Maris and Oostenveld, 2007) on the CD waveforms (time x source vertice x trial matrix) extracted within our ROI for each participant and condition. We defined the post-stimulus as the period between 100 and 250 ms relative to the stimulus onset for each trial and source vertice. This interval, shorter than the full post-stimulus period due to computational constraints, was chosen to include the main components of the visual response (Di Russo et al., 2001). We defined the baseline as the average overtime of the 100 ms prior to stimulus onset for each trial and source vertice. To have a matrix of the baseline (time x source vertice x trial matrix) comparable with the post-stimulus period, the average baseline value for each trial and source vertice was replicated along the time dimension.

In brief, the cluster permutation statistical test starts by clustering individual data samples based on temporal and spatial proximity exhibiting significant t-values (p<.05). The sum of the samples’ t-values belonging to each cluster is submitted to a second-order inference stage. Trial labels (baseline and post-stimulus) are permuted 5000 times to estimate the distribution of maximal cluster-level statistics. A two-tailed Monte-Carlo p-value (p<.05) is then used to identify the significant clusters. While this statistical test identifies the clusters in time and space with significant differences between baseline and post-stimulus, the sign of the t-statistic does not provide information regarding which of the periods has a stronger response in magnitude (Supp. Fig. 1a). For instance, an increase in positive activity from baseline (B) to post-stimulus (P) (*e*.*g*. from 5 to 10 pA.m) produces the same effect as a decrease in negative activity from baseline to post-stimulus (*e*.*g*. from -10 to -5 pA.m). To probe where the response was stronger, we computed the absolute difference between the post-stimulus and baseline period (|P|-|B|, Supp. Fig. 1b), allowing us to distinguish between the two aforementioned cases (|P|-|B|>0 and |P|-|B|<0). The space-time matrix where the post-stimulus activity was higher compared to the baseline is then multiplied by the previously obtained clusters to provide the final matrix (Supp. Fig. 1c). This final matrix contained the clusters in space and time (time x source vertice matrix) that were statistically significant and higher in the post-stimulus period compared to the ongoing activity and was used to mask the evoked CD within the post-stimulus period (time x source vertice matrix).

#### 2.2.4 Measures of inter-hemispheric transfer time

The hypothesis underlying the applied EEG paradigm is that lateralized visual stimuli induce an activity increase in the contralateral hemisphere prior to the ipsilateral one. IHTT can then be estimated at the source-level by taking the latency difference between the two maximal neuronal activations of occipital homologous regions. The maximal neuronal activations are measured on the CD time courses within our ROI.

For each participant and condition, we extracted the CD time courses (of dimension time points x source vertices x number of trials) within our regions of interest in the left and right hemispheres separately. The CDs were averaged across trials (leading to a matrix of dimension time points x source vertices). The absolute value of the resulting CDs were averaged across vertices in each hemisphere, resulting in a time course of CDs (time vector) that was used to estimate IHTT for each participant and condition. For the group IHTT estimation, for each condition, the CD time courses were averaged across subjects to create a group time course.

To estimate IHTT, we first used an automatic peak-picking algorithm (*findpeaks* from MATLAB with minimum peak width of 4 ms) to identify the peak with maximum intensity in the interval between 130 and 220 ms post-stimulus onset in the ipsilateral and contralateral CD time courses. This interval was defined in order to include the two maxima in the CD time courses at the group level. IHTT was calculated by subtracting the latency of the peak associated with the direct path response (contralateral activation – *e*.*g*. right hemisphere for the LVF) from the latency of the peak of the indirect path response (ipsilateral – *e*.*g*. left hemisphere for the LVF). This procedure produced an IHTT estimation based on the reconstructed time course in the source space.

To confirm the origin of the estimated IHTT as the result of a visual-evoked activity, we computed the IHTT from an additional two waveforms obtained from the statistical analysis: the CDs masked with the masking matrix of the significant clusters (Section 2.2.3) and the time course of the number of vertices of the significant clusters of the same masking matrix. Estimating IHTT based on the number of vertices within the significant clusters rather than on the waveforms of activity represents another proxy for the interhemispheric transfer given that the significant activation is expected to increase spatially first in the contralateral and then later in the ipsilateral hemisphere. As previously, we used the same peak-picking algorithm (*findpeaks* from MATLAB) to extract the peak of activity within our ROI in these two waveforms and estimated IHTT by taking the latency difference between the ipsilateral and contralateral peaks.

The consistency of the IHTT estimates obtained from the original CDs, from the CDs masked with the significant clusters, and from the time course of the number of significant vertices was measured with a Pearson’s Correlation Coefficient.

All codes are available on our online repository: https://github.com/DNC-EEG-platform/SingleSubjectIHTTEstimation.

##### 2.2.4.1 Peak selection reliability

To identify the subjects and conditions for which the IHTT estimates were uncertain due to the presence of multiple peaks, we searched in the three types of waveforms all possible maxima falling within 5% of the global maximum, using the *findpeaks* MATLAB command.

##### 2.2.4.2 Within-session and between-session repeatability

The within-session repeatability of the IHTT estimates obtained on the original CDs waveforms was assessed by subdividing the trials from the first session of each participant into two non-overlapping random subsets and estimating the IHTT on each of the subsets. The degree of the agreement of IHTT estimates between the two subsets was assessed with Pearson’s Correlation Coefficient across participants. To evaluate the magnitude of the within-session IHTT variability we computed the absolute difference between the IHTT estimates from the two subsets of trials.

To measure the repeatability of the IHTT estimates between sessions, we performed a second EEG recording session on three of the participants and estimated IHTT on the original CDs waveforms. The degree of the agreement of IHTT estimates between sessions across participants was assessed with Pearson’s Correlation Coefficient. To evaluate the magnitude of the between-session IHTT variability we computed the mean absolute difference between the IHTT estimates across participants.

### 2.3 Magnetic Resonance Imaging-Based Estimation of the G-Ratio

#### 2.3.1 Data acquisition

The analysis conducted is as reported in our previous work (Oliveira et al., 2022). In brief, all MRI scans were collected on a 3T MRI system (Magnetom Prisma; Siemens Medical Systems, Erlangen, Germany). The MRI acquisition included: a 3D structural T1-weighted Magnetization-Prepared Rapid Gradient-Echo (MPRAGE) image with a 1 mm^3^ isotropic voxel size; a relaxometry protocol that was used to compute maps of the Magnetization Transfer (MTsat) with 1 mm^3^ isotropic voxel size; and a diffusion protocol along 15, 30, and 60 diffusion directions with b = 650/1,000/2,000 s/mm^2^, respectively, with 2 mm^2^ isotropic resolution.

#### 2.3.2 Data processing

A full description of the data analysis can be found in Oliveira et al., 2022. Freesurfer (Fischl, 2012) was used to delineate the occipital ROI (defined in Section 2.2.3). mrTrix (Tournier et al., 2019) was used to delineate the tract that connected the occipital ROI of the two brain hemispheres and crossed the corpus callosum. MTsat maps were calculated with the hMRI toolbox (Tabelow et al., 2019) from the images obtained with the relaxometry protocol. Eddy current correction in the diffusion data were corrected with FSL (Andersson and Sotiropoulos, 2016) and image distortions were corrected using the fieldmap toolbox of SPM (Hutton et al., 2002). The NODDI model (Zhang et al., 2012) was fitted to the preprocessed diffusion data, leading to maps of the isotropic diffusion (*v*_*iso*_) and intracellular (*v*_*ic*_) compartments volume fractions. Finally, maps of gMRI were calculated from the MTsat maps and from *v*_*iso*_ and *v*_*ic*_ maps (Stikov et al., 2015): 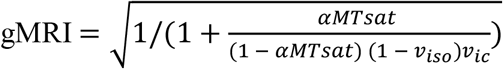, with *α* = 0.23 (Slater et al., 2019). The MRI-measured g-ratio maps were sampled along the white matter transcallosal tract delineated with diffusion MRI tractography.

### 2.4 Axonal morphology analysis

Estimates of axonal conduction velocity were computed by dividing the length of the transcallosal white matter tract in each dataset by the corresponding estimate of IHTT from the original CDs. Subsequently, the measures of gMRI and conduction velocity were used to estimate morphological properties of axons in the transcallosal white matter tract using the model proposed in (Oliveira et al., 2022). The axonal morphological properties included:

- the scale parameter *θ* of the distribution of axonal radius a white matter tract. *θ* represents the tail of the distribution, *i*.*e*. the fraction of large axons within a tract
- the scaling factor *β* of the dependence of the axonal g-ratio on axonal radius. *β* represents the amplitude of the change in axonal g-ratio with axonal radius.

The model parameters α and *M* were set to 0.14 and 0.30 µm respectively, in agreement with histological studies (Tomasi et al., 2012; Liewald et al., 2014).

The specificity of the *θ* and *β* estimates computed from each dataset was assessed by comparing intra-subject variability and inter-subject variability. Intra-subject variability was estimated as the mean difference of the *θ* and *β* estimates between sessions. Inter-subject variability was estimated as the standard deviation of the *θ* and *β* estimates across the 14 datasets of the first session.

## 3 Results

### 3.1 Group IHTT estimation

Group-averaged CD waveforms exhibits a sharp increase compared to the ongoing activity starting at approximately 115 ms post-stimulus onset (Fig. 1). This increase is first observed in the hemisphere contralateral to the visual stimulation – right hemisphere (red line in Fig. 1) in the case of an LVF stimulation – and later in the ipsilateral hemisphere – left hemisphere (blue line in Fig. 1) in the case of an LVF stimulation. These results confirm our assumptions on the expected pattern of neuronal activations, *i*.*e*. a response of the hemisphere directly stimulated (direct pathway) occurring earlier than the opposite one (indirect pathway). The group-averaged estimates of IHTT was 11 and 19 ms for the LVF and RVF respectively.

**Fig 1.**
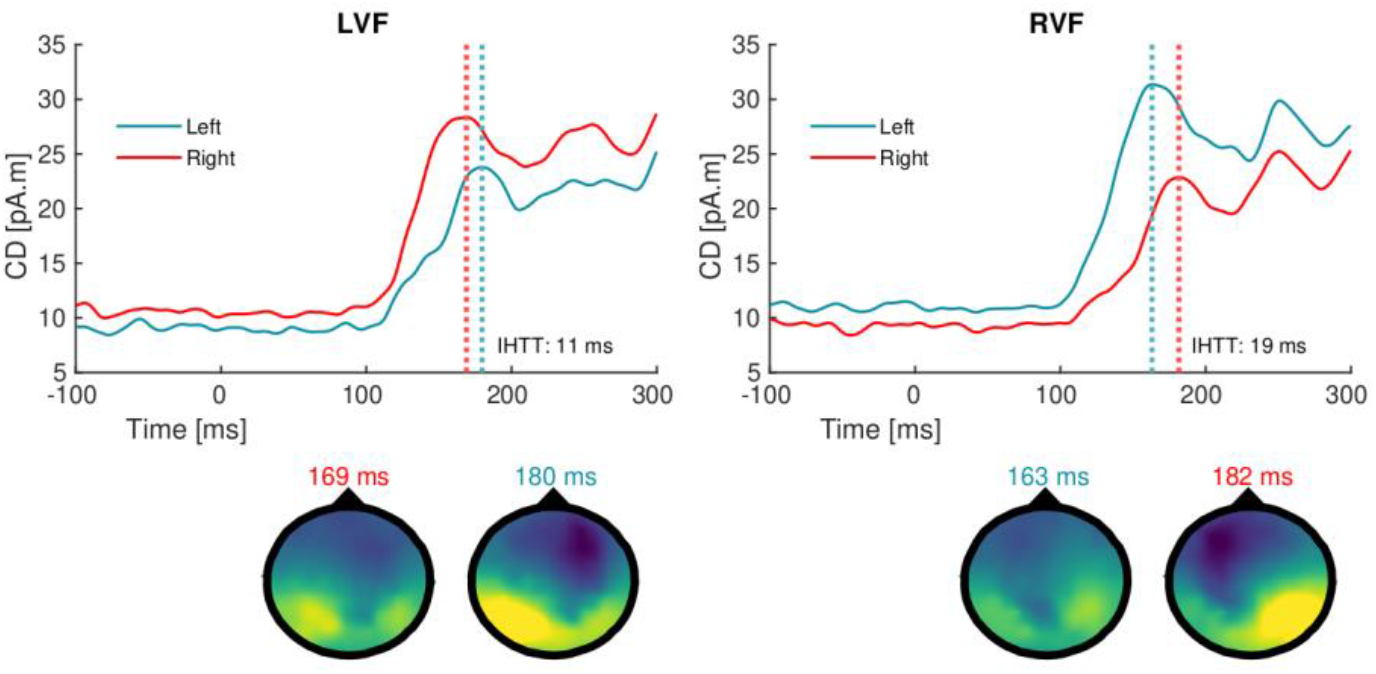
Group averaged CD for the LVF (left) and RVF (right) conditions. Stimulus onset took place at t = 0ms. The CDs inside the occipital ROI on the left and right hemispheres are depicted in blue and red lines, respectively. The peaks of activation in each hemisphere are shown by the vertical dashed lines. Voltage topographies maps are shown for the time points corresponding to the identified peaks.

### 3.2 Subject-specific IHTT estimation

Similarly to the group CD waveforms (Fig. 1), the time course of the individual original CD waveforms typically exhibit a first peak in the contralateral hemisphere, followed by a peak in the ipsilateral hemisphere (solid lines in Fig. 2 a,b). The same peak latencies are identified by the CD waveforms masked with the results of the cluster permutation statistical analysis, indicating that these peaks lie within the period of the stimulus-evoked activity (dashed lines in Fig. 2 a,b). The latency of the peaks on the CD waveforms are consistent with the latencies of the ERP component N1 (∼120-210 ms) as observed in the GFP waveforms of the corresponding ERPs (Fig. 2 c,d). The cluster permutation identifies clusters of activation that increase in size with time; the clusters on the contralateral and ipsilateral hemispheres show a time-lag between them (Fig. 2 e,f). The latencies of the maximum number of activation vertices in each hemisphere approximately correspond to the latencies of the peaks seen on the CD waveforms. The results for the remaining subjects can be found in Supp. Fig. 2.

**Fig 2.**
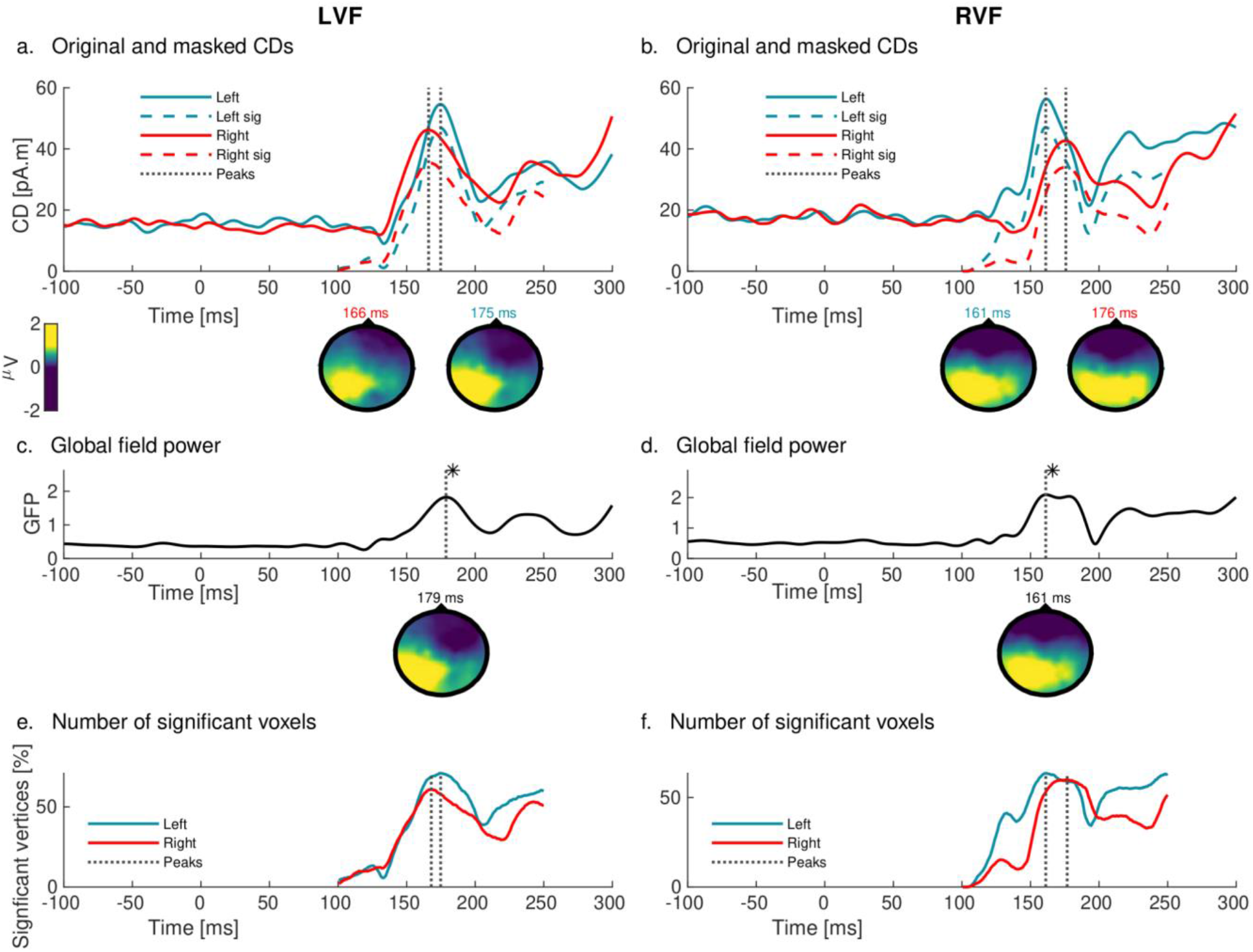
Summary of the IHTT estimation in an exemplar participant a, b) CD waveforms for one example subject for both visual conditions (LVF and RVF). The original CDs inside the occipital ROI on the left and right hemispheres are depicted with blue and red solid lines, respectively. The masked CDs in the same ROI on the left and right hemispheres are shown in the blue and red dashed lines, respectively. The vertical dashed lines identify the selected peaks in each hemisphere on the original data. The voltage topographies correspond to those identified peaks. c,d) Global field power time course and topographies corresponding to the main components. The * identifies the component closest to the first peak identified in a, b). e, f) Number of significant voxels (percentage) in the post-stimulus period. Left and right hemispheres’ timecourses are shown with blue and red lines, respectively. Vertical lines correspond to peaks. Time 0 ms identifies the onset of the stimulus for all the graphs.

Table 1 shows for each subject and condition the IHTT estimated from the original CDs, the CDs waveforms masked with the significant clusters, and the time course of the number of vertices inside the significant clusters. A positive IHTT reflects a transfer in the direction predicted anatomically, where the response of the hemisphere directly stimulated (direct pathway) occurs earlier than the opposite one (indirect pathway). For the LVF, the estimated IHTT was in the direction predicted anatomically in 12 subjects out of 14 from the original CDs and the masked CDs, and in 10 subjects out of 14 from the significant number of vertices. For the RVF, IHTT was in the direction predicted anatomically in 9 subjects out of 14 from the original CDs and the significant number of vertices and 8 subjects out of 14 from the masked CDs. The mean IHTT across conditions from the original CDs ranged from 8 to 29 ms across participants (with a mean of 19 ms), excluding one participant with a negative IHTT of -25 ms.

**Table 1.**
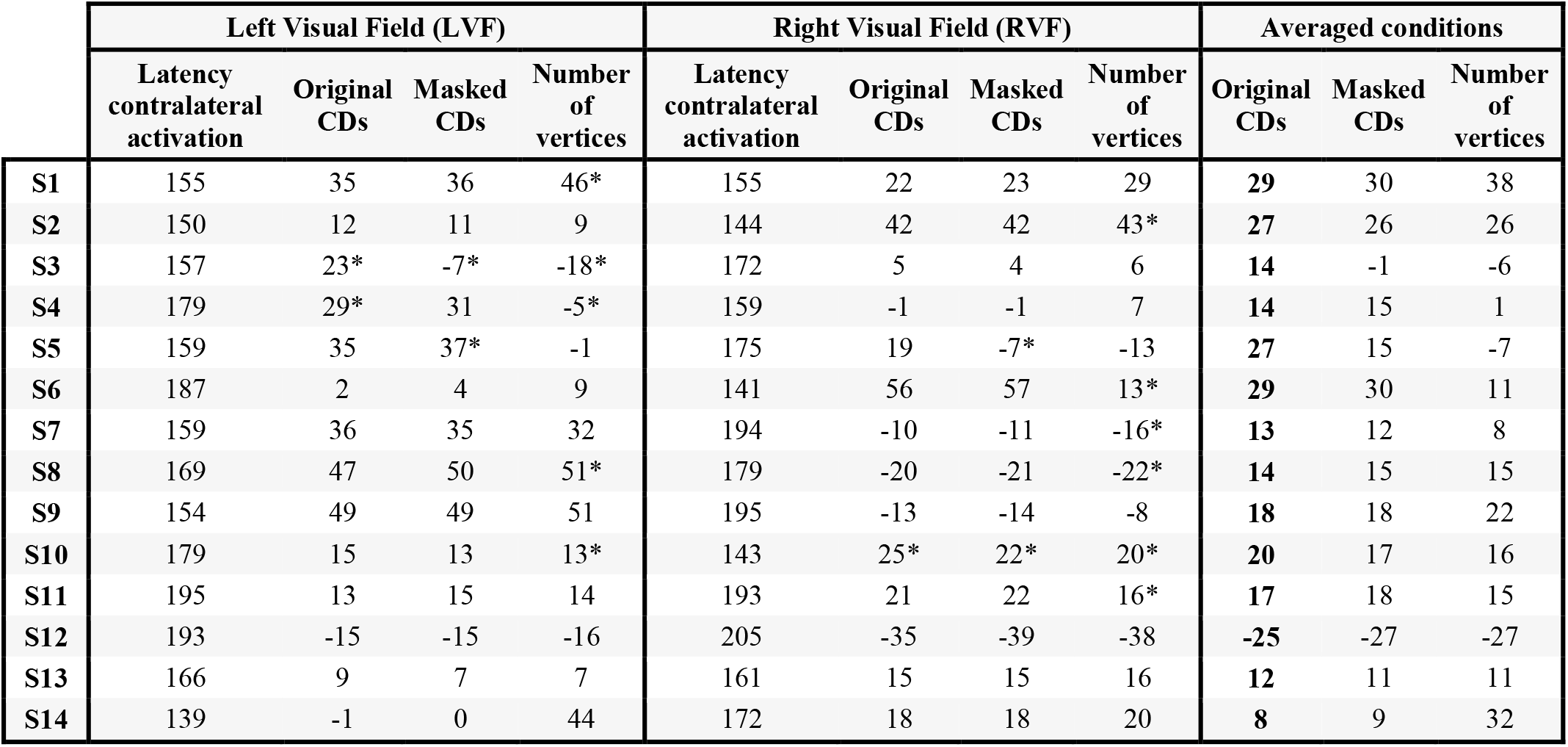
Overview of the IHTT estimation at the single-subject level. Latency contralateral activation refers to the contralateral peak latency as measured from the original CDs. IHTT estimates for the LVF and RVF with the three metrics for the 14 subjects and average across conditions. Highlighted in bold are the values of the IHTT used for the morphological estimates. * identifies the cases where the waveforms contained multiple peaks with amplitude values within 5% of the global maximum and therefore considered less reliable than the values without *. Time is always shown in ms.

The correlation coefficient of the IHTT estimates from the original CDs and masked CDs was 0.95, from the original CDs and number of voxels was 0.68 and between the masked CDs and number of voxels was 0.76. In all three cases, the correlation coefficient was significant with p<.001.

### 3.3 Peak selection reliability

To identify unreliable IHTT estimates due to the presence of multiple peaks, we searched for a second peak with an amplitude within 5% of the global maximum (* in Table 1). For the LVF, unreliable IHTT estimates were found in 2 subjects from the original CDs or masked CDs and in 5 subjects from the significant number of vertices. Similarly for the RVF, unreliable IHTT estimates were found in 1 subject from the original CDs, in 2 subjects from the masked CDs, and in 6 subjects from the significant number of vertices. The IHTT estimated from the first and second highest peaks is detailed in Supp. Table 1. The mean absolute difference between the IHTT measured on the first and the second highest maximum of neuronal activity is 32 ms for the LVF and 26 ms for the RVF.

By considering the second-highest amplitude maximum on the CD waveforms, the disagreement between the metrics is often mitigated. This can be observed in the LVF of S3, where the IHTT obtained on original data is positive (23 ms, Fig. 3a). The IHTT obtained on the masked data is negative and thus anatomically unplausible when obtained with the absolute maximum (−7 ms, Fig. 3b) but when calculated with the second highest maximum is positive and in-line with the one obtained from the original data (24 ms, Fig. 3c).

**Fig 3.**
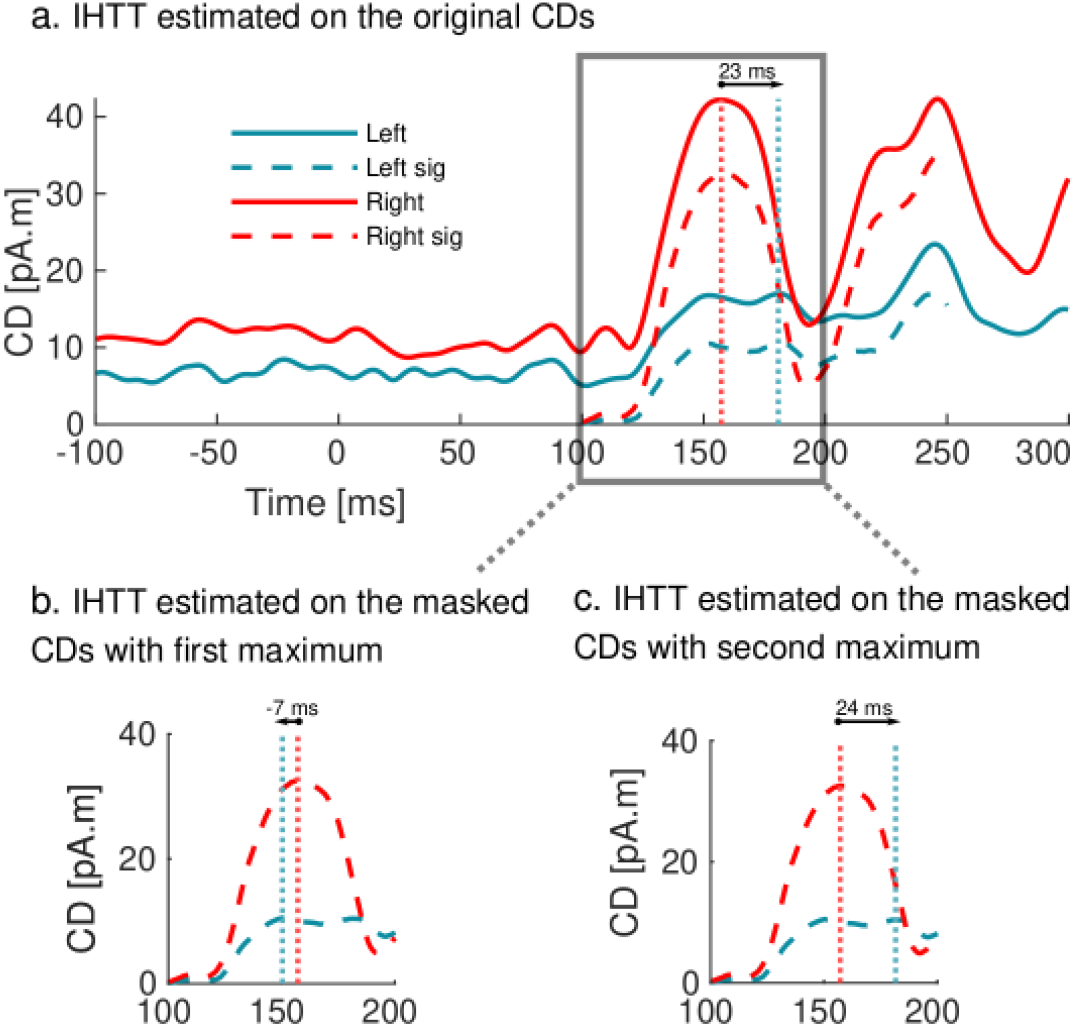
Example of the presence of multiple peaks with amplitude within 5% of the global maximum. a) Original CDs inside the occipital ROI on the left and right hemispheres are depicted with blue and red solid lines, respectively. The masked CDs in the same ROI on the left and right hemispheres are shown with blue and red dashed lines, respectively. Time 0 ms identifies the onset of the stimulus. The vertical dashed lines identify the selected peaks on the original CDs. The estimated IHTT is 23 ms. b) Zoom in on the masked CDs of (a). The vertical blue dashed line identifies the global maximum of the left hemisphere. The estimated IHTT is -7 ms. c) Zoom in on the masked CDs of (a). The vertical blue dashed line identifies the second-highest amplitude peak in the left hemisphere. The estimated IHTT is 24 ms, in-line with the estimated IHTT on the original CDs shown in (a).

### 3.4 IHTT within-session and between-session repeatability

For the within-session repeatability, the IHTT estimates from the first subset of trials showed moderate repeatability with the ones from the second subset of trials (r=0.68, p<.001, Fig. 4a). The mean IHTT absolute difference across the two subsets was ∼11 ms with some differences reaching up to 63 ms. The IHTT estimates obtained from each subset (circles in Fig. 4b) differ from the IHTT value obtained from all the trials (horizontal lines in Fig. 4b): the absolute differences between the mean IHTT across subsets and the IHTT obtained from all the trials are on average 6 ms for the LVF and 5 ms for the RVF.

**Fig 4.**
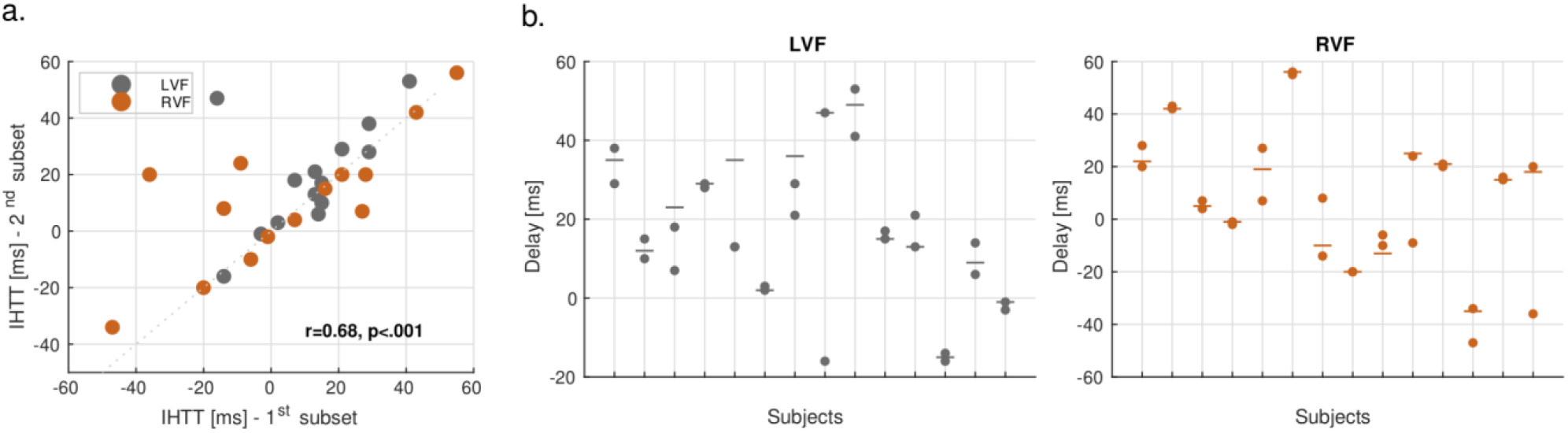
a) IHTT was obtained from two non-overlapping subsets of the trials for each subject for both conditions for the first session. There is a significant correlation (r = 0.68, p<.001) between the two subsets. b) Relation between the IHTT obtained for the two non-overlapping subsets of the trials (circles) and the IHTT obtained with all the trials (horizontal lines) for each participant and condition. The mean absolute difference between the latter and the mean IHTT across subsets was on average 6 ms for the LVF and 5 ms for the RVF.

For the between-session repeatability, three participants were scanned in a second session. For the LVF, the IHTT was 12, 23, and 36 ms in the first session and 12, 17, and 20 ms in the second session. As for the RVF, the IHTT was 42, 5, and -10 ms in the first session and 5, -1, and -7 ms in the second session. The absolute difference in IHTT across sessions can reach up to 37 ms, with an average value of 11 ms. Nonetheless, visually the CD waveforms show a high level of correspondence (Fig. 5).

**Fig 5.**
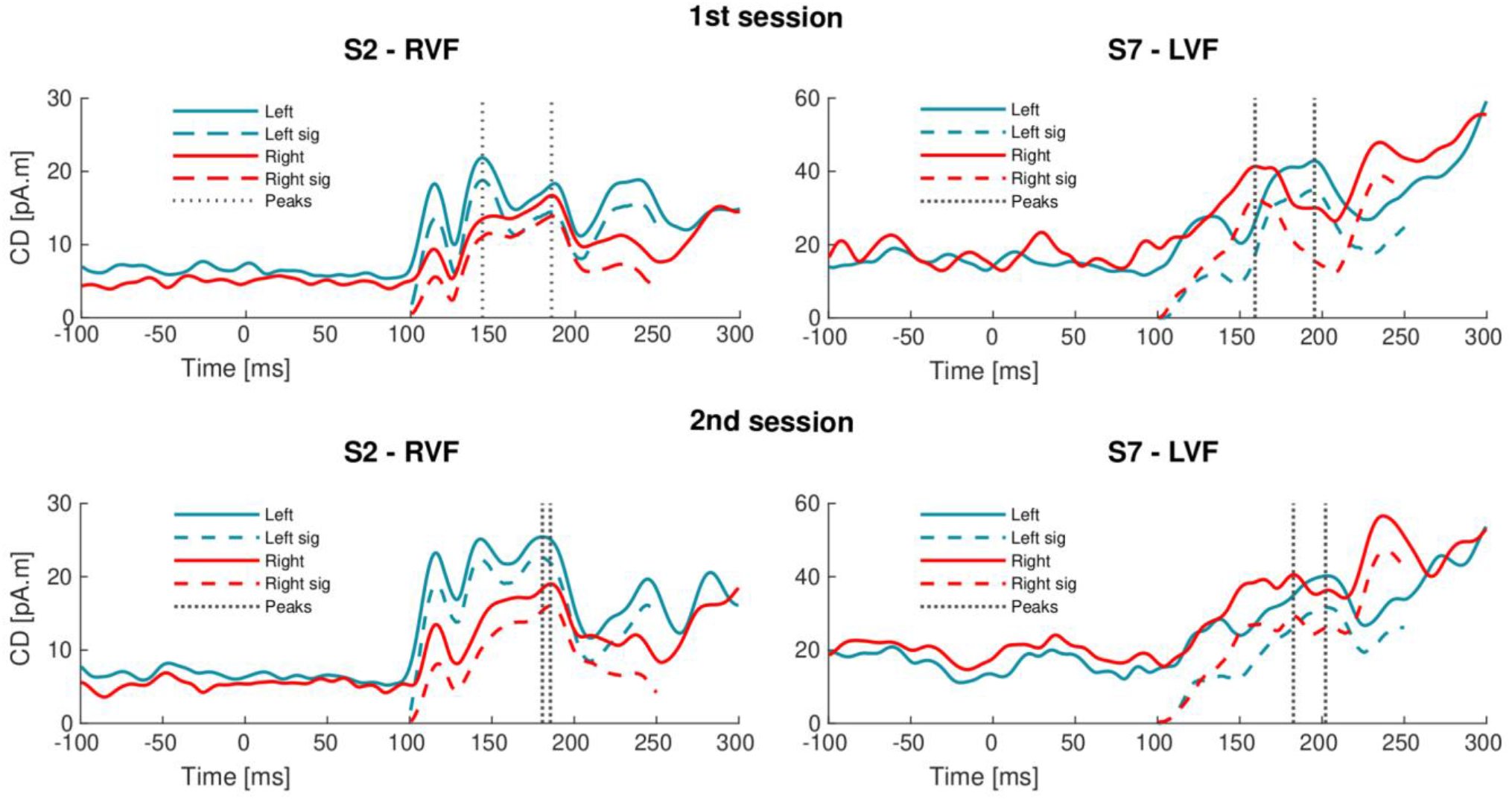
CD waveforms for the two sessions for subjects S2 (RVF) and S7 (LVF). The original CDs inside the occipital ROI on the left and right hemispheres are depicted in the blue and red solid lines, respectively. The masked CDs in the same ROI on the left and right hemispheres are shown in the blue and red dashed lines, respectively. The vertical dashed lines identify the selected peaks in each hemisphere on the original data. Time 0 ms identifies the onset of the stimulus for all the graphs.

### 3.5 Estimation of axonal morphology from the MRI and EEG data

The conduction velocities ranged from 4.7 to 18.5 m/s (excluding one negative conduction velocity, Table 2).

**Table 2.** Final IHTT estimates averaged across conditions for the 14 subjects (ms). Length of the white matter tract (mm). Conduction velocity along the white matter tract (m/s). Estimated θ (*μ*m) and β (*μm*^−*α*^) for each subject.

| | IHTT estimated<br>from original CDs<br>[ms] | Length [mm] | Velocity [m/s] | $\theta$ [$\mu\text{m}$] | $\beta$ [$\mu\text{m}^{-\alpha}$] |
| --- | --- | --- | --- | --- | --- |
| S1 | 29 | 136.5 | 4.7 | 0.00 | 0.81 |
| S2 | 27 | 134.6 | 5.0 | 0.02 | 0.81 |
| S3 | 14 | 127.7 | 9.1 | 0.26 | 0.71 |
| S4 | 14 | 133.6 | 9.5 | 0.27 | 0.69 |
| S5 | 27 | 149.0 | 5.5 | 0.03 | 0.77 |
| S6 | 29 | 147.3 | 5.1 | 0.01 | 0.79 |
| S7 | 13 | 143.9 | 11.1 | 0.38 | 0.69 |
| S8 | 14 | 136.3 | 9.7 | 0.29 | 0.70 |
| S9 | 18 | 141.6 | 7.9 | 0.18 | 0.72 |
| S10 | 20 | 135.4 | 6.8 | 0.11 | 0.74 |
| S11 | 17 | 155.8 | 9.2 | 0.25 | 0.70 |
| S12 | -25 | 155.6 | -6.2 | - | - |
| S13 | 12 | 154.2 | 12.9 | 0.46 | 0.65 |
| S14 | 8 | 147.8 | 18.5 | 0.79 | 0.62 |

These conduction velocities and the gMRI values sampled along the occipital transcallosal tract were used to estimate the parameter *θ* of the axonal radius distribution and the parameter *β* of the change in axonal g-ratio with axonal radius (Table 2). The average *θ* across subjects was 0.23 *μ*m and ranged from 0.00 to 0.79 *μ*m (Table 2). Subject S1 is the only one with a *θ* value of almost zero (< 0.01 *μ*m). The average *β* across subjects was 0.72 *μm*^−*α*^ ranging from 0.62 to 0.81 *μm*^−*α*^ (Table 2). The inter-subject variability in the model estimates, given by the standard deviation of the estimated parameters, is 0.22 *μ*m and 0.06 *μm*^−*α*^ for *θ* and *β* respectively. A qualitative impression of the variability of the axonal radius distribution and axonal fiber myelination across the cohort is provided in Fig. 6a and Fig. 6b, respectively.

**Fig 6.**
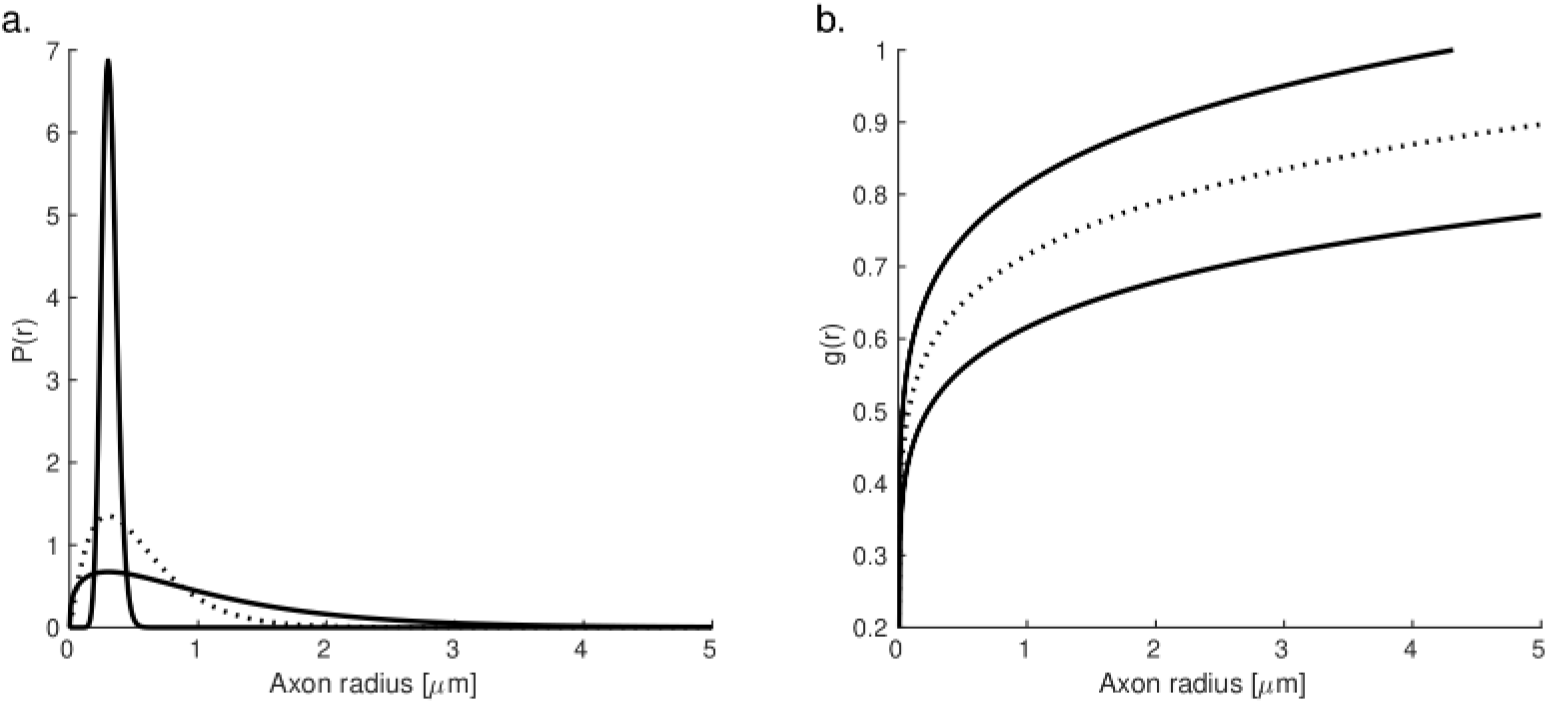
Minimum, maximum (solid lines), and average (dashed line) model estimates across subjects: a) Axonal radius distribution P(r) and b) axonal g-ratio g(r).

The between-sessions variability of the axonal morphology estimates was estimated from the data in a second session, acquired from 3 participants. The mean difference in *θ* and *β* between sessions was 0.57 *μ*m and 0.11 *μm*^−*α*^ respectively, larger than inter-subject variability. To better understand the origin of this high variability, we plotted the values of the *θ* (Fig. 7a) and *β* (Fig. 7b) estimates obtained from each subject within the 2-dimensional space of possible combinations of conduction velocities and gMRI. This result shows that the conduction velocity is the main direction of change for the *θ* and *β* estimates and that the repeatability of conduction velocity estimates is the primary determinant of the repeatability of axonal morphology estimates.

**Fig 7.**
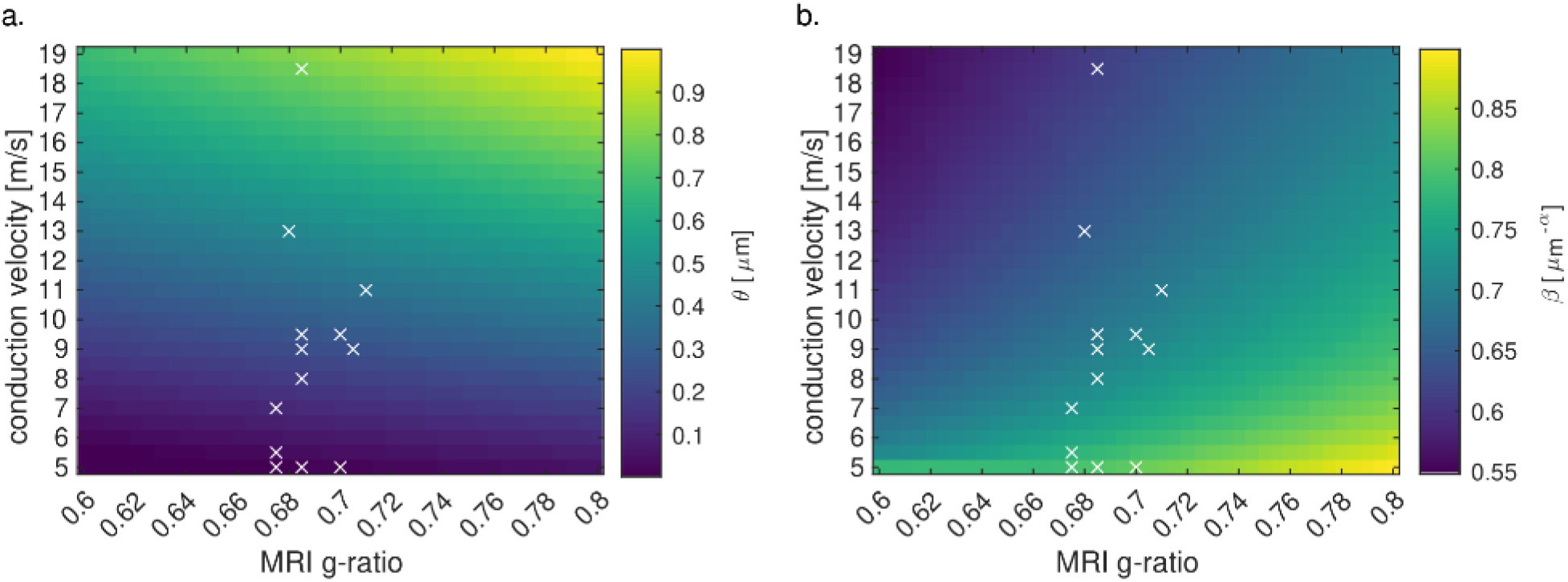
Range of the model parameters θ (a) and β (b) computed from combinations of simulated in vivo MRI g-ratio and conduction velocities. The crosses indicate the subject-specific MRI g-ratio and conduction velocities. Conduction velocities are in the main direction of change in θ and β estimates.

Supplementary analyses show that a 10% bias in conduction velocity leads to an average bias in *θ* of about 22%, within the plausible range of in-vivo data values (Supp. Fig. 3); and an average bias in *β* of approximately 2%. These results highlight the importance of accurate conduction velocity estimates for the model estimates.

## 4 Discussion

In this work, we proposed a novel framework to estimate the IHTT at the single-subject level from EEG data, based on a data-driven evaluation of the maximal peak of neural response to visual stimuli with minimal *a priori* constraints. The resulting subject-specific estimates of IHTT were used to estimate morphological features of axons within the occipital transcallosal white matter tract, using a recently introduced biophysical model (Oliveira et al., 2022).

The proposed framework relies on the estimation of the neural activity in the source space. This avoids the *ad hoc* selection of electrodes and ERP components at the electrode level and overcomes ambiguities related to the undetermined relation between voltage measurements at the scalp and those of the underlying and neurophysiologically interpretable electric fields (Nunez and Srinivasan, 2009). The subject-specific IHTT estimates are in agreement with the literature (average = 18.6 ms), except for one negative value. From the values of the IHTT, the scale parameter of the axonal radius distribution (*θ*), which represents the fraction of large axons within a white matter tract, was estimated as 0.23 *μ*m on average across subjects. The amplitude of the change in axonal g-ratio with axonal radius (*β*) is 0.72 *μm*^−*α*^ on average across subjects, in-line with the literature.

### 4.1 IHTT measures

The IHTT estimates obtained from the original CDs at the group level are consistent with previous literature (Saron and Davidson, 1989; Friedrich et al., 2017; Chaumillon et al., 2018). Moreover, faster right-to-left transfer (*i*.*e*. LVF, 11 ms) than left-to-right transfer (*i*.*e*. RVF, 19 ms) is consistent with previous findings in right-hand dominated cohorts (Marzi et al., 1991; Martin et al., 2007; Moes et al., 2007; Whitford et al., 2011; Chaumillon et al., 2018).

As for the subject-specific estimates, the mean IHTT across the two visual conditions (LVF and RVF) ranged from 8 to 29 ms across participants – excluding one participant with a negative IHTT of -25 ms (Table 1). These results are in agreement with the few electrode-based studies that report subject-specific IHTT values (Westerhausen et al., 2006; Friedrich et al., 2017; Chaumillon et al., 2018). Our range of estimated IHTT is also in-line with the delays estimated with intracranial recordings from whole-brain cortical-cortical connections (Lemarechal et al., 2022).

The reliability of the subject-specific measures of IHTT was assessed by estimating the consistency of the IHTT estimates computed from the original CDs, masked CDs, and number of significant vertices. The correlation coefficients between two pairs of metrics were above 0.68 and significant (p<.001) in all cases. The overall agreement between the three metrics confirms the origin of the estimated IHTT on the original CDs as the result of a stimulus-elicited activity.

Analysis of the repeatability of the IHTT estimates within-session (Fig. 4a), conducted from non-overlapping subsets of trials for each participant, showed moderate repeatability (r=0.68, p<.001). The average variability in IHTT within-session was 11 ms, approximately half of the mean IHTT across subjects. The between-session repeatability, analyzed on a subgroup of 3 participants, shows a weak linear relationship between the IHTT estimated on two different sessions. The average variability between sessions was 11 ms. This is comparable to within-session variability despite contributions from environmental or contextual factors such as activation strength and attentional resource allocations.

The variability of the IHTT estimates between-session was also comparable to the differences in IHTT estimates between subjects (7 ms), suggesting that the IHTT estimates exhibit a low level of specificity to each participant. Nevertheless, the pattern of neuronal activity is qualitatively reproducible across sessions (Fig. 5). This result suggests that peaks of maximal activation might be insufficient to estimate the IHTT. Other approaches, such as transfer entropy (Vicente et al., 2011) or mutual information (Ince et al., 2017) could be considered as future alternatives at the expense of a high number of parameters and assumptions. Confidence intervals on the IHTT values for each subject could also be estimated with a bootstrapping approach, by sampling the delays from the subject’s trial data with replacement (Cichy et al., 2014).

### 4.2 Subject-specific neuronal activity

At the single-subject level, only one subject (S12) did not fulfill our expected patterns of activity - *i*.*e*. increase in activation in the contralateral hemisphere prior to the ipsilateral hemisphere - and thus has negative IHTT in both visual conditions. The reason for this remains unclear. We postulate that the simple process that we assume to describe the visual transfer (peak in one hemisphere and then transfer to the other hemisphere) might be too simplistic. For instance, multiple transfers might occur within a short period (Deslauriers-Gauthier et al., 2019). Also, undeniably, rather than a single IHTT value, there exists a distribution of delays due to the underlying axonal radius distribution (Caminiti et al., 2013).

While the peaks of maximal neuronal response are very evident in the group CD waveforms (Fig. 1), the same is not always true for the individual data. Variability in the latencies and topographies of the neuronal response, and in the cortical anatomy across subjects has been extensively reported (Foxe and Simpson, 2002; Proverbio et al., 2007; Baumgartner et al., 2018). Hence, understanding the specific dynamics of each individual data (Supp. Fig. 2) is fundamental when analyzing the data-driven IHTT estimates. In particular, the presence of multiple peaks with analogous amplitudes on the waveforms hinders the IHTT estimation. If we take such cases into account (Suppl. Table 1), the overall agreement between IHTT metrics is even more evident. These results support the existence of a subject-specific neuronal response, the difficulty being in the definition of a robust metric to extract IHTT from the CD time courses that is consistent across subjects and requires a minimal set of assumptions.

The presence of multiple peaks in at least two of the metrics was observed in three cases (the LVF of S3 (Fig. 3a), RVF of S10, and LVF of S4). For these participants, two possible IHTT estimates exist for all three metrics, with only one of the two options being anatomically plausible (*i*.*e*. positive IHTT). For example, the LVF IHTT of S3 may be 23, 24, and 23 ms or -5, -7, and -18 ms from the original CDs, masked CDs, and number of significant vertices, and no objective metric exists to guide the choice of a maximum. Because our expected pattern of activity is a contralateral peak occurring before the ipsilateral one (positive IHTT), we could potentially impose an extra constraint on the positivity of the IHTT. However, that would diverge from a data-driven approach with minimal constraints which we are proposing here.

In summary, we use a simple assumption for the pattern of interhemispheric transfer, with minimal constraints to obtain single-subject IHTT estimates. We showed that with our approach, estimating the subject-specific IHTT based on the original CDs is associated with some level of uncertainty due to the presence of peaks with similar amplitudes in only 3 out of 28 cases (Section 3.3). In any case, for these 3 cases, the maximum peak, which is the one we chose by default, is the one providing an IHTT in the direction predicted anatomically. The IHTT based on the original CDs is also supported by the other two metrics and the agreement of the IHTT estimates with the literature is compelling.

### 4.3 Subject-specific axonal morphologic estimates

The parameter *θ* represents the fraction of axons with large radius within a white matter tract, and ranged from 0.00 to 0.79 *μ*m (Fig. 6a). The non-plausible *θ* value of 0.00 *μ*m is associated with one of the subjects with the highest IHTT (S1: 29 ms); the next lower *θ* value is 0.01 *μ*m. For the maximum value of *θ* (0.79 *μ*m), axons above 2 *μ*m represent <14% of the total fiber count, in agreement with histological studies showing maximal axonal radius in the human brain of ∼1.5–3 µm (Aboitiz et al., 1992; Caminiti et al., 2009; Liewald et al., 2014). The mean radius corresponding to *θ* values of 0.01 *μ*m and 0.79 *μ*m are 0.31 *μ*m and 1.09 *μ*m respectively, approximately inline with previous estimates from histological studies (0.62 *μ*m; Caminiti et al., 2009). The parameter *β*, which represents the amplitude of the change in axonal g-ratio with axonal radius, ranged from 0.62 to 0.81 *μm*^−*α*^ (Fig. 6b).

The cornerstone of the proposed white matter model is that the structural (MRI g-ratio) and functional (conduction velocity) measures originate from the same white matter tract: the MRI g-ratio values are sampled along this tract and the IHTT estimates are divided by the tract’s length used to compute the conduction velocity. The extent of evoked activity from primary to high-order visual areas in response to visual stimulation may imply the involvement of multiple white matter tracts for interhemispheric transfer. As these tracts remain undefined, the tract length was chosen as the mean length of the streamlines that connect the occipital regions across both hemispheres. This tract is 12 mm (8%) smaller than the tract that connects the primary and secondary visual cortices used in our previous study (Oliveira et al., 2022). Differences in gMRI values between these tracts were minimal (<1%).

It is worth pointing out that, unlike Oliveira et al., 2022, conduction velocities were measured from an average of the two experimental conditions (LVF and RVF). This allows for a more representative characterization of the white matter tract, independent of the direction of propagation of neurons.

### 4.4 Future directions

This study focuses on the single-subject measurement of the IHTT, which is essential to the assessment of axonal morphology in-vivo in separate individuals. The consistency of the axonal radius and myelination estimates with the histological literature highlights the long-term potential of this technique. However, we found that the variability of the morphological estimates between repetitions was in the order of – or larger than – the difference of these estimates between subjects. Our results suggest that the measurement of the IHTT is the primary source of variability in the estimates of axonal morphology. Increasing the reproducibility of the IHTT estimates may be achieved by considering other source reconstruction methods, alternative metrics for IHTT estimation from the CD time course, or different EEG paradigms with a focus on other tracts. Alternative techniques, such as the newly proposed approach for single-subject and single-tract IHTT estimation, based on resting-state EEG data (Sorrentino et al., 2022) should also be considered. Comparison of EEG-based IHTT estimates with intracranial recordings of local field potentials and histological measures are essential to ensure their validity as in-vivo measures of axonal conduction velocity.

## 5 Conclusions

This work represents the first attempt to estimate morphological properties of white matter axons from combined MRI and EEG data acquired in-vivo in individual subjects. In particular, we propose a new data-driven framework with minimal *a priori* constraints that allows the single-subject measurement of the IHTT from EEG data, based on the maximal peak of neural response upon visual stimulation. This framework provides evidence that the IHTT estimates are the result of activity elicited by the visual stimulus. The estimated IHTT values are in the reported range of electrode-based and intracranial EEG studies (Saron and Davidson, 1989; Friedrich et al., 2017; Chaumillon et al., 2018; Lemarechal et al., 2022). The MRI data and EEG-based measures of the IHTT were used to estimate morphological properties of white matter axons. The agreement of the estimates of axonal radius and myelination with the histological literature highlights the long-term potential of this technique. However, these morphological estimates showed a high level of variability that arises primarily from the estimation of the IHTT. Increasing the repeatability of the IHTT estimates and comparison with alternative measures of conduction velocity are essential future steps toward the measurement of axonal morphological properties from in-vivo data.

## 8 Acknowledgments

The MRI data were acquired on the MRI platform of the Clinical Neuroscience Department, Lausanne University Hospital. We thank all technicians involved in the data collection and researchers Andria Pelentritou, Giulia Di Domenicantonio and Quentin Raynaud for their invaluable help with EEG and MRI recordings and insightful discussions.

## 10 Data Availability Statement

The data that support the findings of this study will be available online upon publication of this article.

