## Supplementary for "Single-subject EEG measurement of interhemispheric transfer-time for the in-vivo estimation of axonal morphology"

### Supplementary Material

Supplementary Fig. 1:

Schematic representation of the analysis steps for estimating the evoked current density (CD) in response to the visual stimuli that are statistically significant during the post-stimulus with respect to the baseline period. a. First, cluster permutation was performed on the CD waveforms (time x source vertice x trial matrix) extracted within our region of interest for each participant and condition by comparing the baseline with the post-stimulus period. This process identifies the clusters in time and space with significant differences between baseline and post-stimulus (orange squares). This analysis provides information on when and where the post-stimulus CD is different from the baseline CD, irrespective of whether the post-stimulus absolute amplitude is higher or lower than the baseline one. b. In the second step, we took the absolute difference between the post-stimulus and the baseline in the average CD responses (time x source vertice matrix) to identify the post-stimulus activity that is higher in magnitude compared to the baseline (orange squares). c. The multiplication of the results of a. and b. leads to the final matrix containing the clusters in space and time (orange squares) that are statistically significant and higher in the post-stimulus period compared to the ongoing activity.

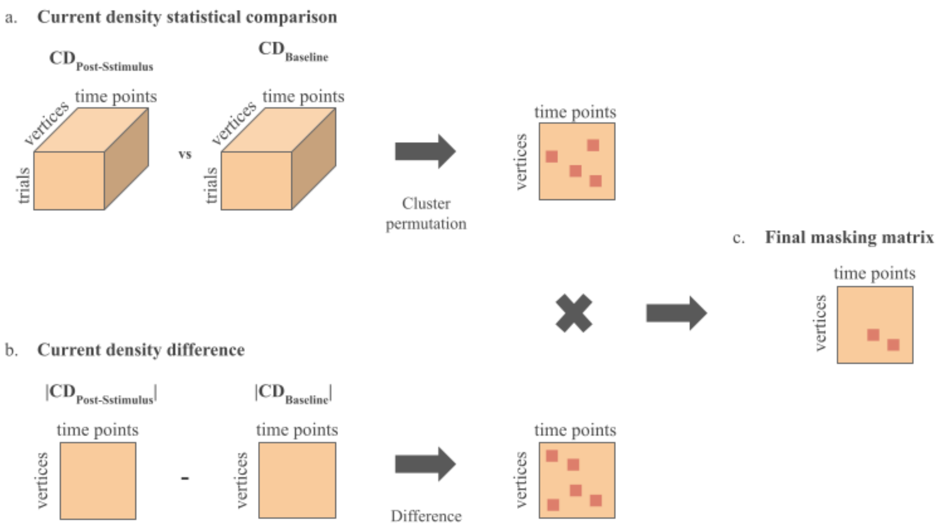

Supplementary Fig. 2:

CD waveforms for all subjects subject for both visual conditions (LVF and RVF). The original CDs inside the occipital ROI on the left and right hemispheres are depicted in the blue and red solid lines, respectively. The masked CDs in the same ROI on the left and right hemispheres are shown in the blue and red dashed lines, respectively. The vertical dashed lines identify the selected peaks in each hemisphere on the original data. The voltage topographies correspond to those identified peaks. c,d) Global field power timecourse and topographies corresponding to the main components. The * identifies the component closest to the first peak identified in a, b). e, f) Number of significant voxels (percentage) in the post-stimulus period. Left and right hemispheres' timecourses are shown in the blue and red lines, respectively. Vertical lines correspond to peaks. Time 0 ms identifies the onset of the stimulus for all the graphs. Similar to Fig. 2 of the main article.

S1

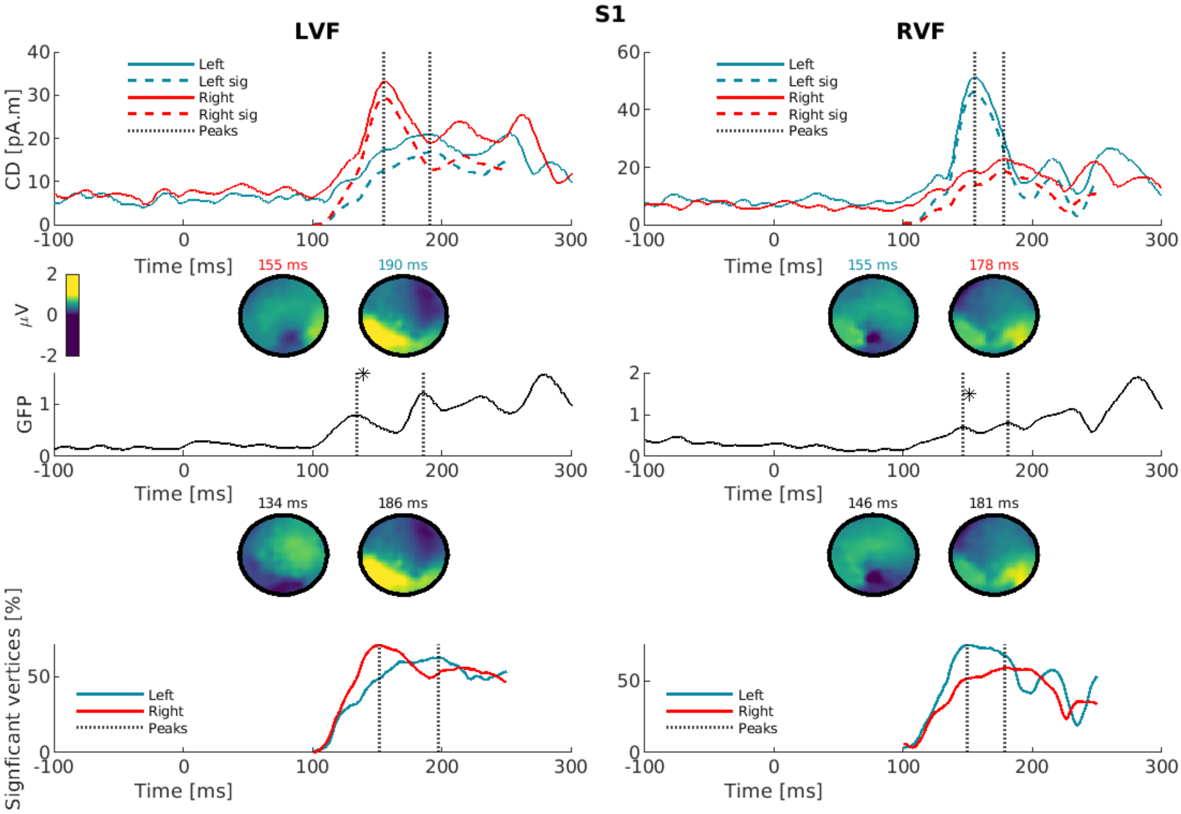
LVF
RVF
CD [pA.m]
Left
Left sig
Right
Right sig
Peaks
Time [ms]
155 ms
190 ms
155 ms
178 ms
μV
GFP
134 ms
186 ms
146 ms
181 ms
Signficant vertices [%]

S2

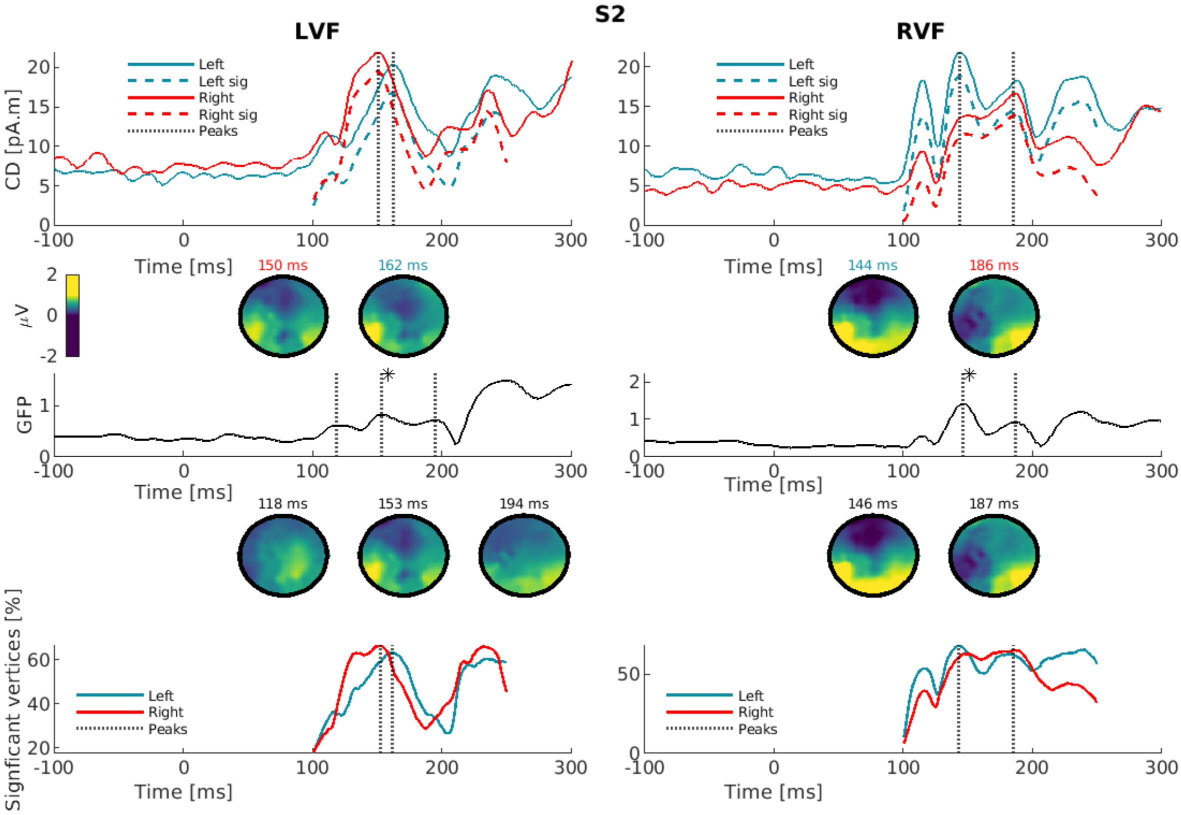
LVF
RVF
CD [pA.m]
Left
Left sig
Right
Right sig
Peaks
Time [ms]
150 ms
162 ms
144 ms
186 ms
μV
GFP
118 ms
153 ms
194 ms
146 ms
187 ms
Signficant vertices [%]

**S3**

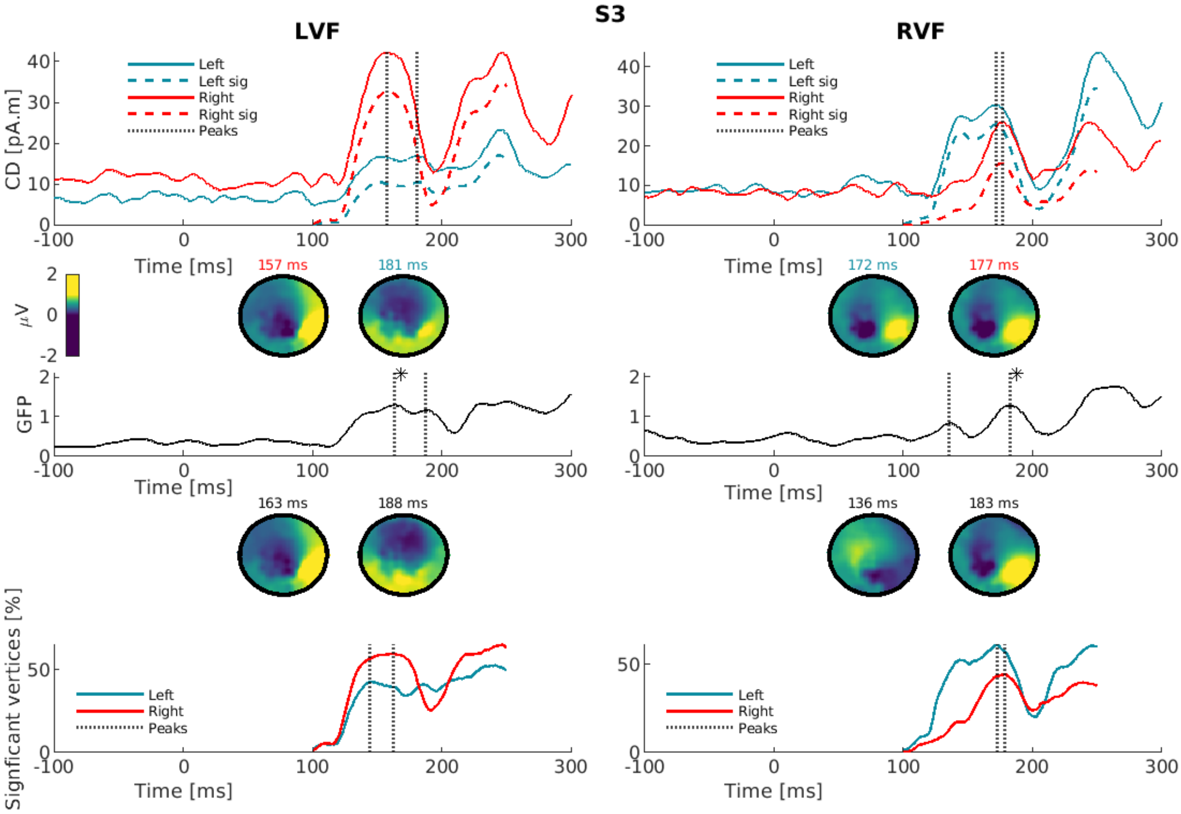
LVF
RVF
Left
Left sig
Right
Right sig
Peaks
CD [pA.m]
Time [ms]
157 ms
181 ms
172 ms
177 ms
µV
GFP
163 ms
188 ms
136 ms
183 ms
Signficant vertices [%]

**S4**

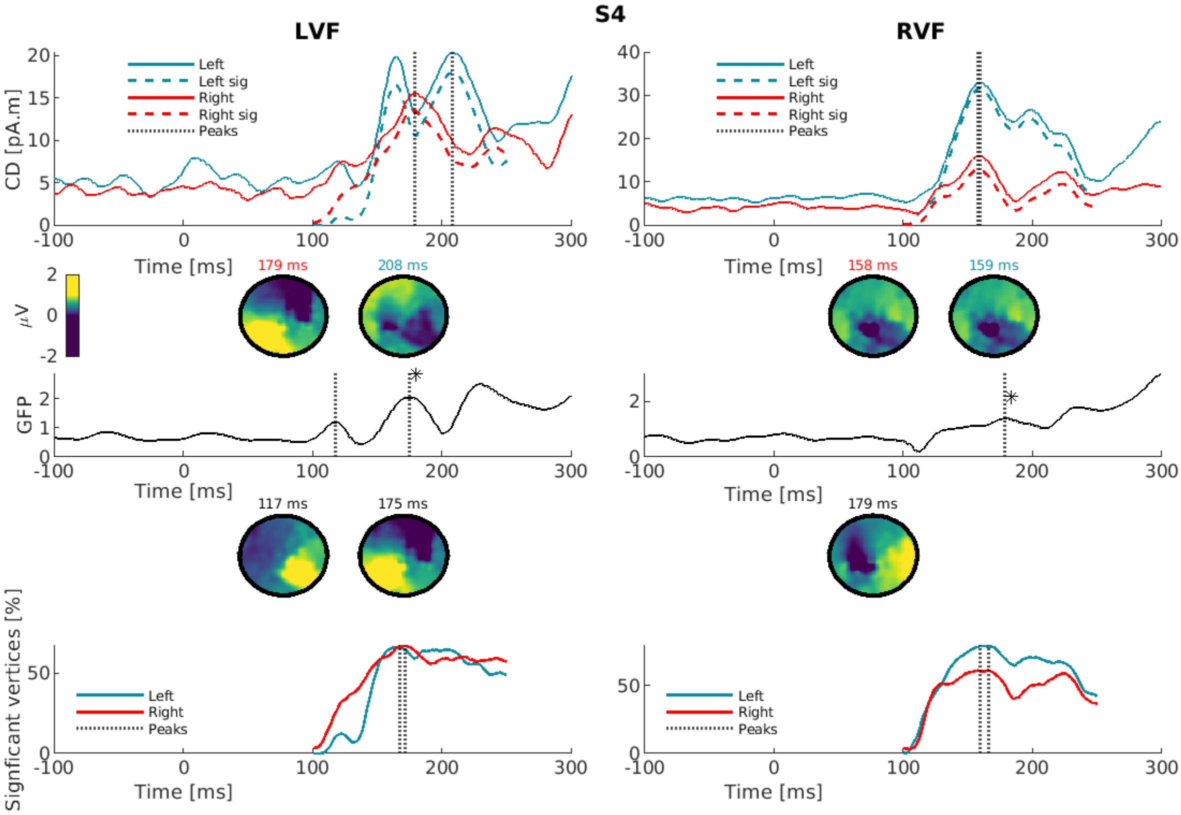
LVF
RVF
Left
Left sig
Right
Right sig
Peaks
CD [pA.m]
Time [ms]
179 ms
208 ms
158 ms
159 ms
µV
GFP
117 ms
175 ms
179 ms
Signficant vertices [%]

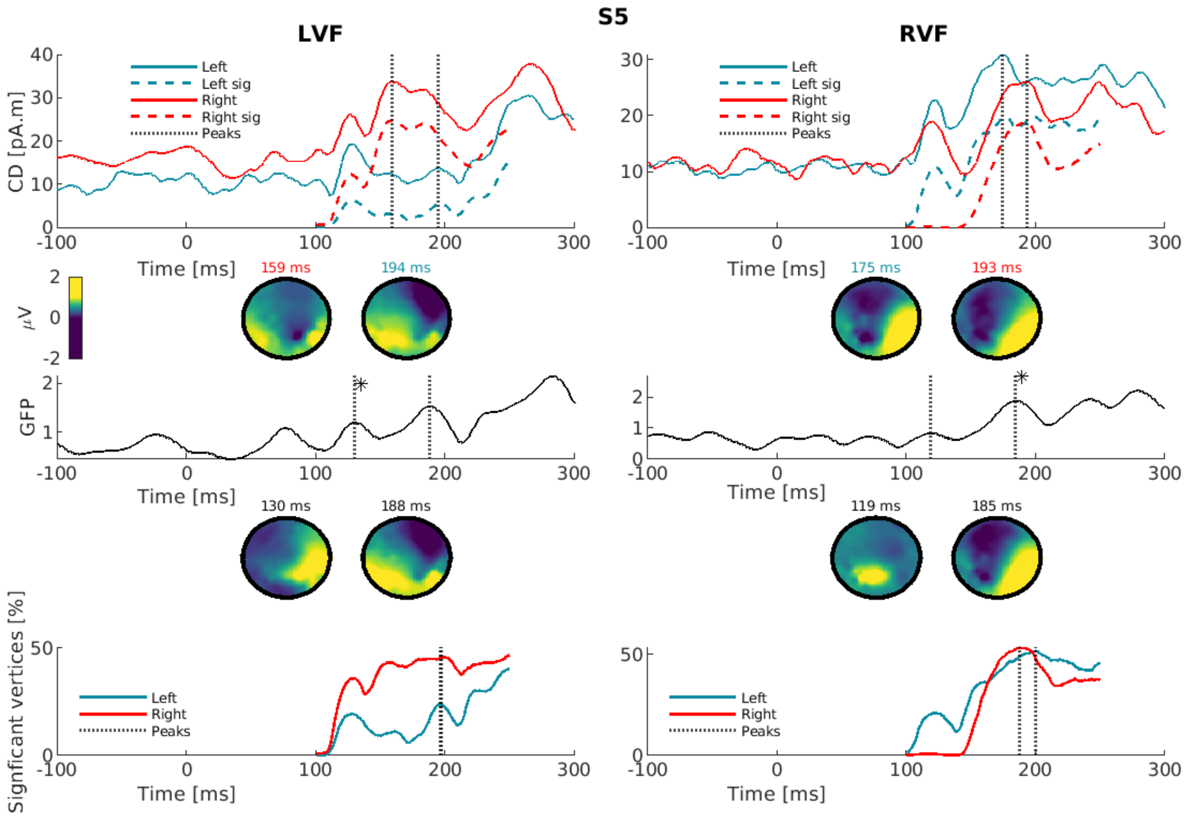
S5
LVF
RVF
Left
Left sig
Right
Right sig
Peaks
CD [pA.m]
Time [ms]
159 ms
194 ms
175 ms
193 ms
μV
GFP
130 ms
188 ms
119 ms
185 ms
Signficant vertices [%]

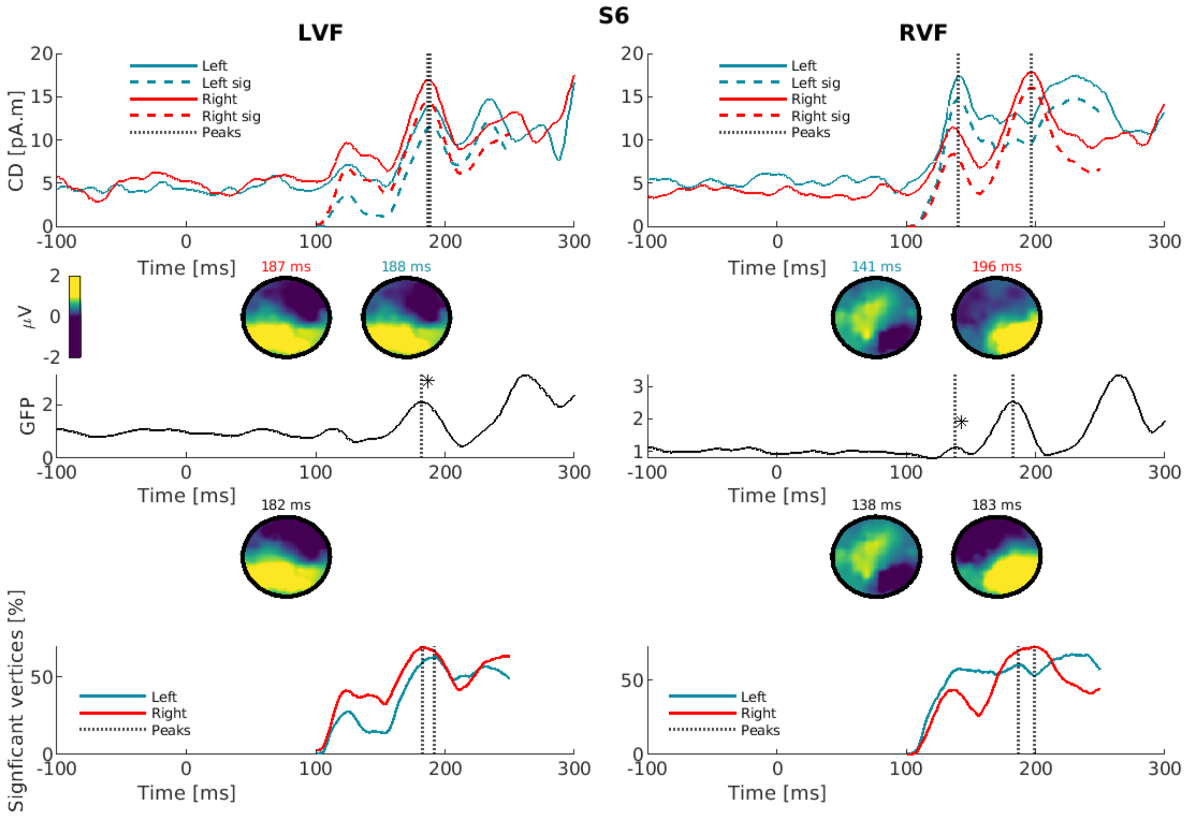
S6
LVF
RVF
Left
Left sig
Right
Right sig
Peaks
CD [pA.m]
Time [ms]
187 ms
188 ms
141 ms
196 ms
μV
GFP
182 ms
138 ms
183 ms
Signficant vertices [%]

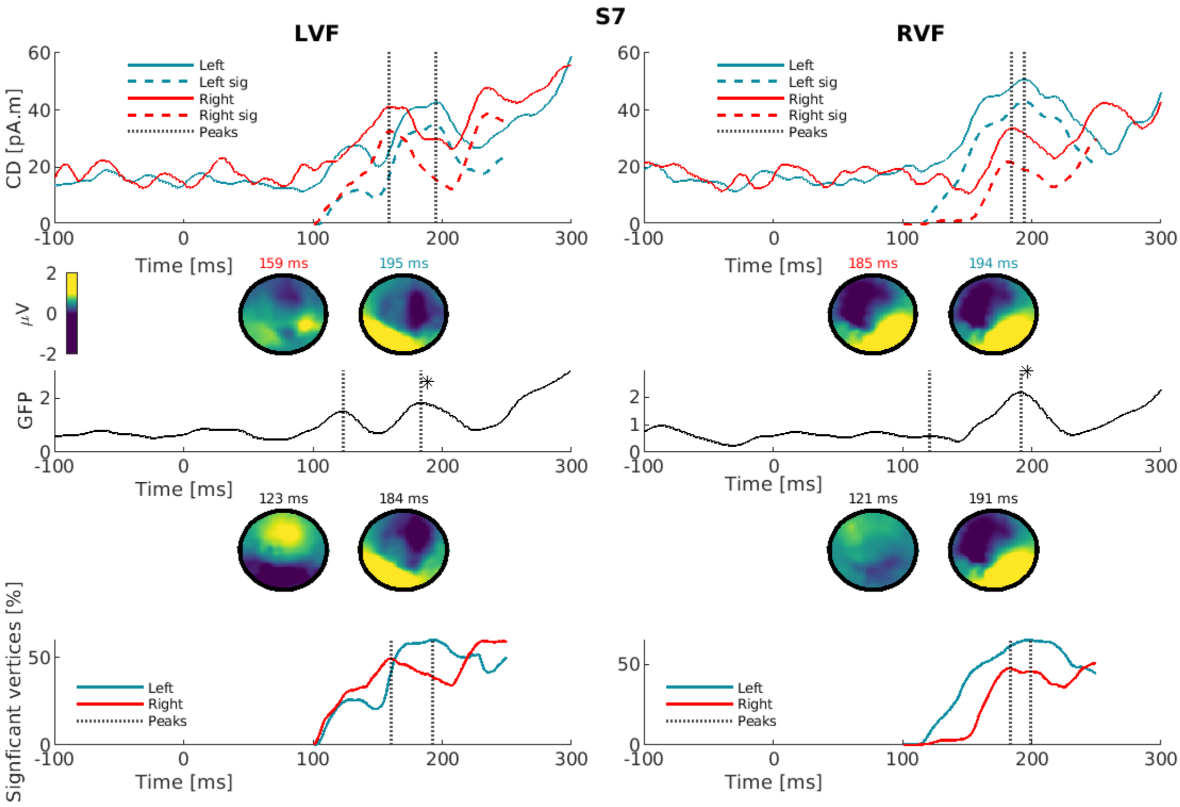
S7
LVF
RVF
Left
Left sig
Right
Right sig
Peaks
CD [pA.m]
Time [ms]
159 ms
195 ms
185 ms
194 ms
μV
GFP
123 ms
184 ms
121 ms
191 ms
Signficant vertices [%]

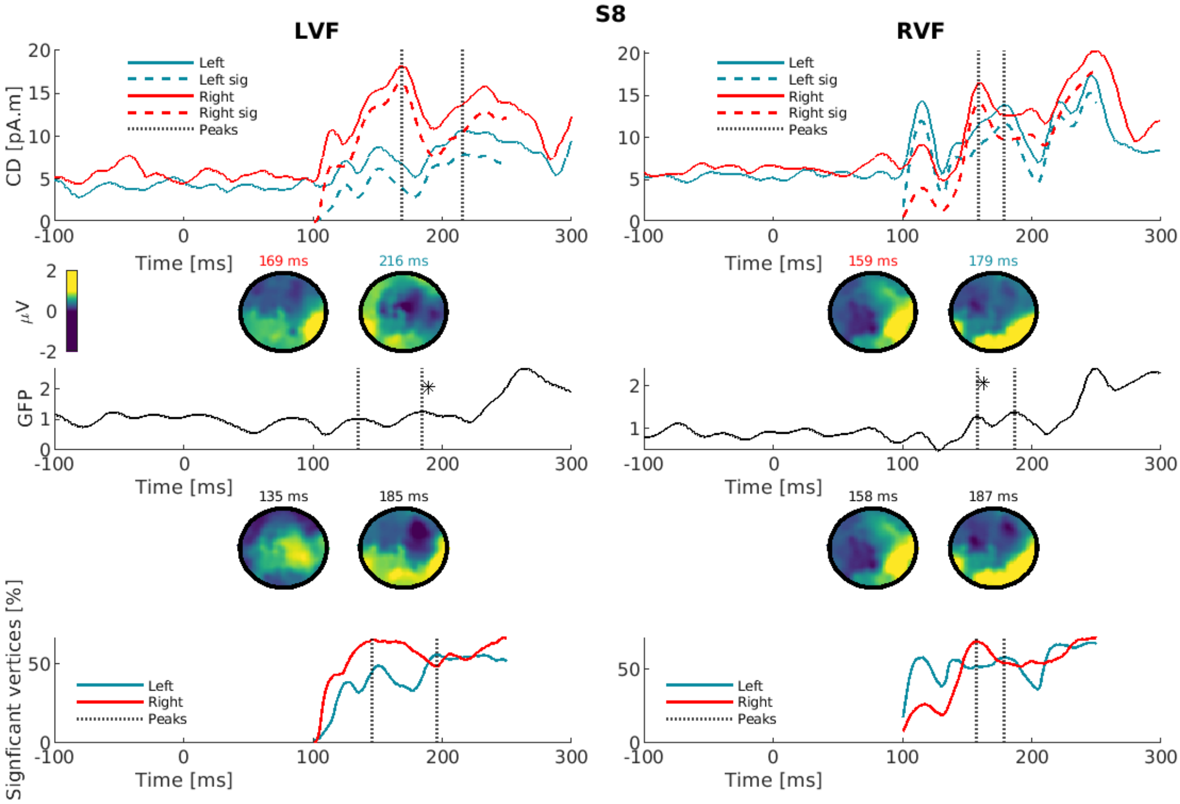
S8
LVF
RVF
Left
Left sig
Right
Right sig
Peaks
CD [pA.m]
Time [ms]
169 ms
216 ms
159 ms
179 ms
μV
GFP
135 ms
185 ms
158 ms
187 ms
Signficant vertices [%]

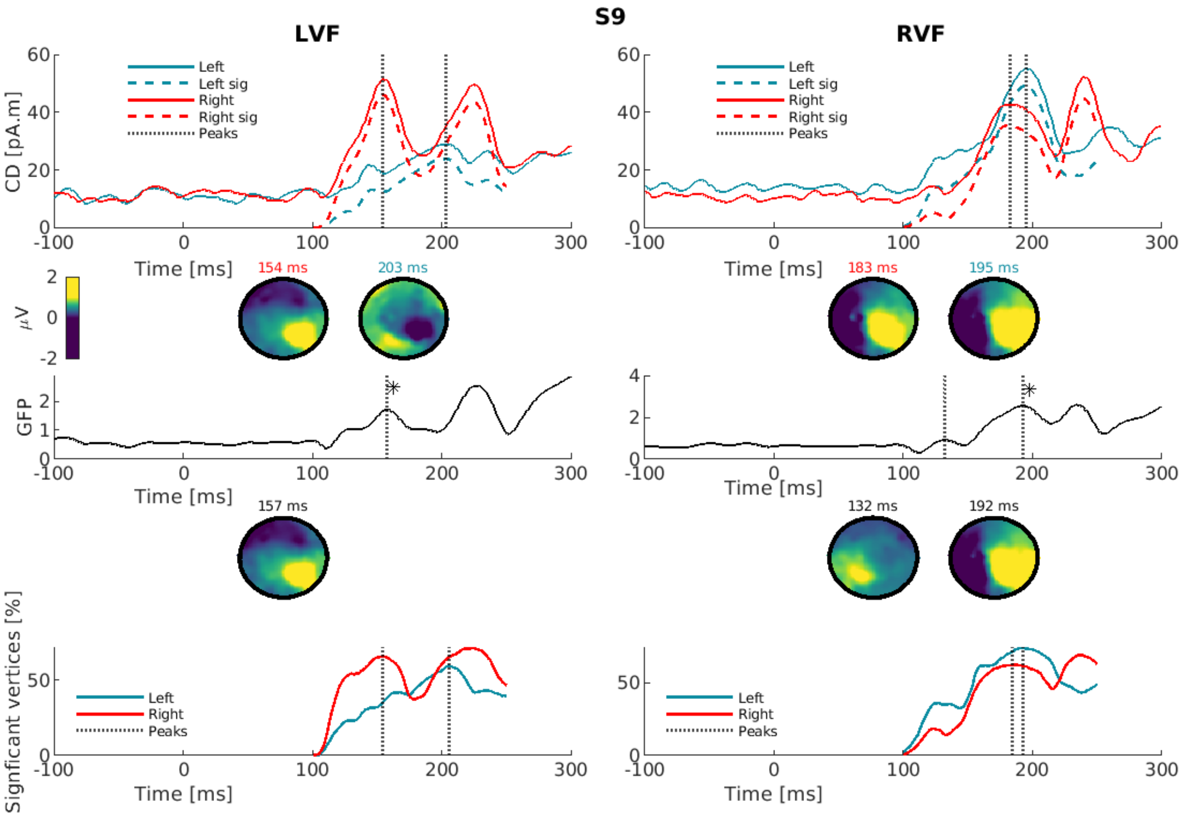
S9
LVF
RVF
CD [pA.m]
Time [ms]
Left
Left sig
Right
Right sig
Peaks
154 ms
203 ms
183 ms
195 ms
μV
GFP
157 ms
132 ms
192 ms
Signficant vertices [%]

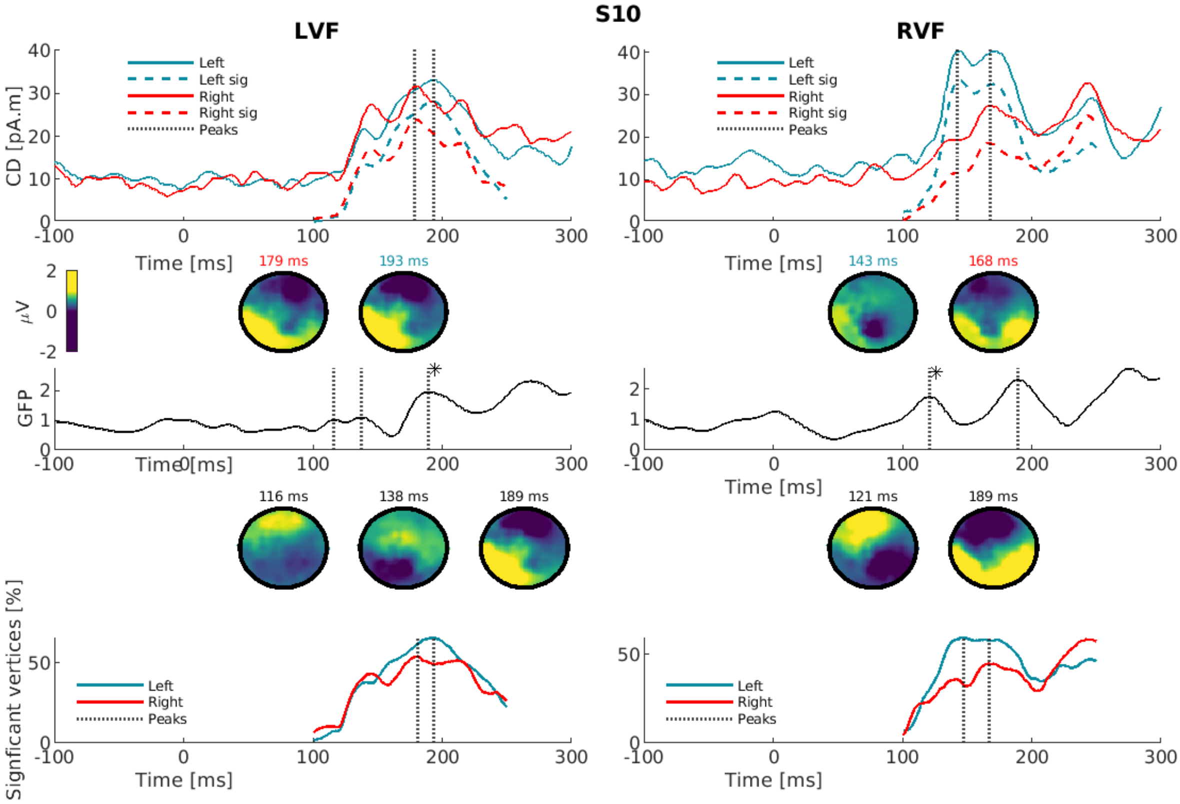
S10
LVF
RVF
CD [pA.m]
Time [ms]
Left
Left sig
Right
Right sig
Peaks
179 ms
193 ms
143 ms
168 ms
μV
GFP
116 ms
138 ms
189 ms
121 ms
189 ms
Signficant vertices [%]

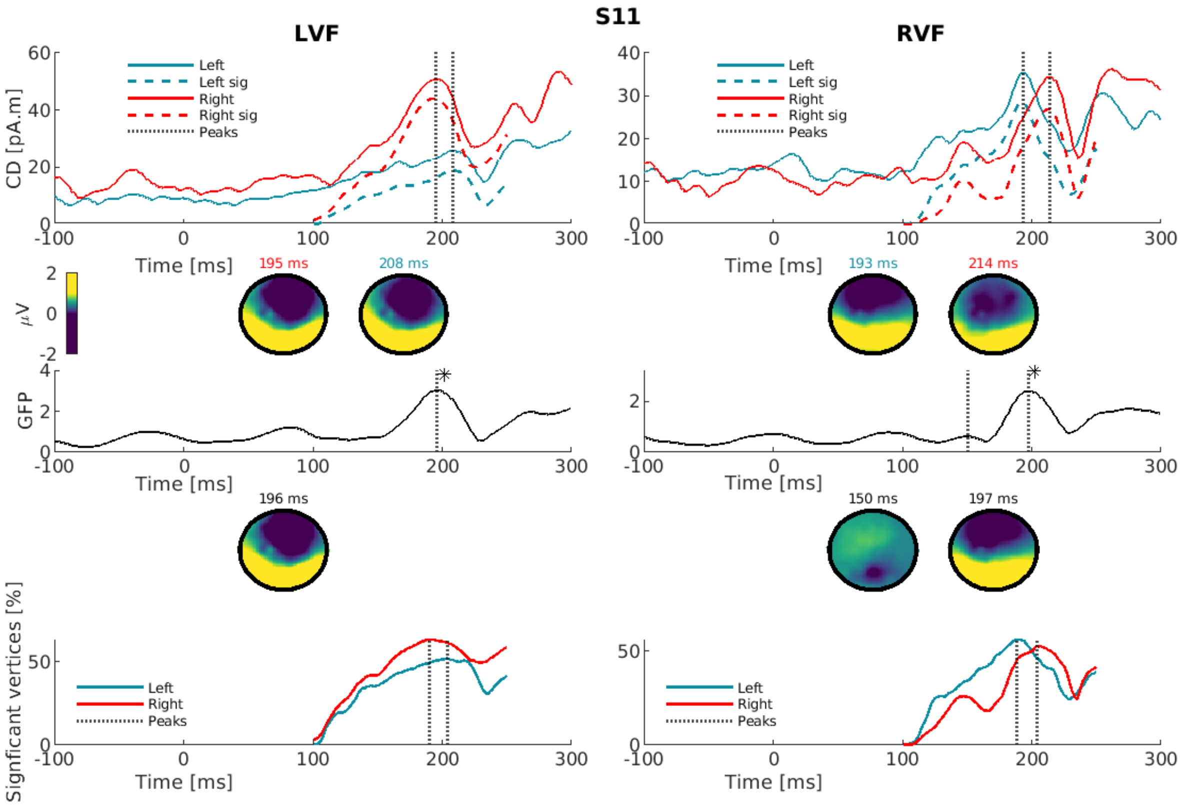
S11
LVF
RVF
CD [pA.m]
Left
Left sig
Right
Right sig
Peaks
Time [ms]
195 ms
208 ms
193 ms
214 ms
μV
GFP
196 ms
150 ms
197 ms
Signficant vertices [%]

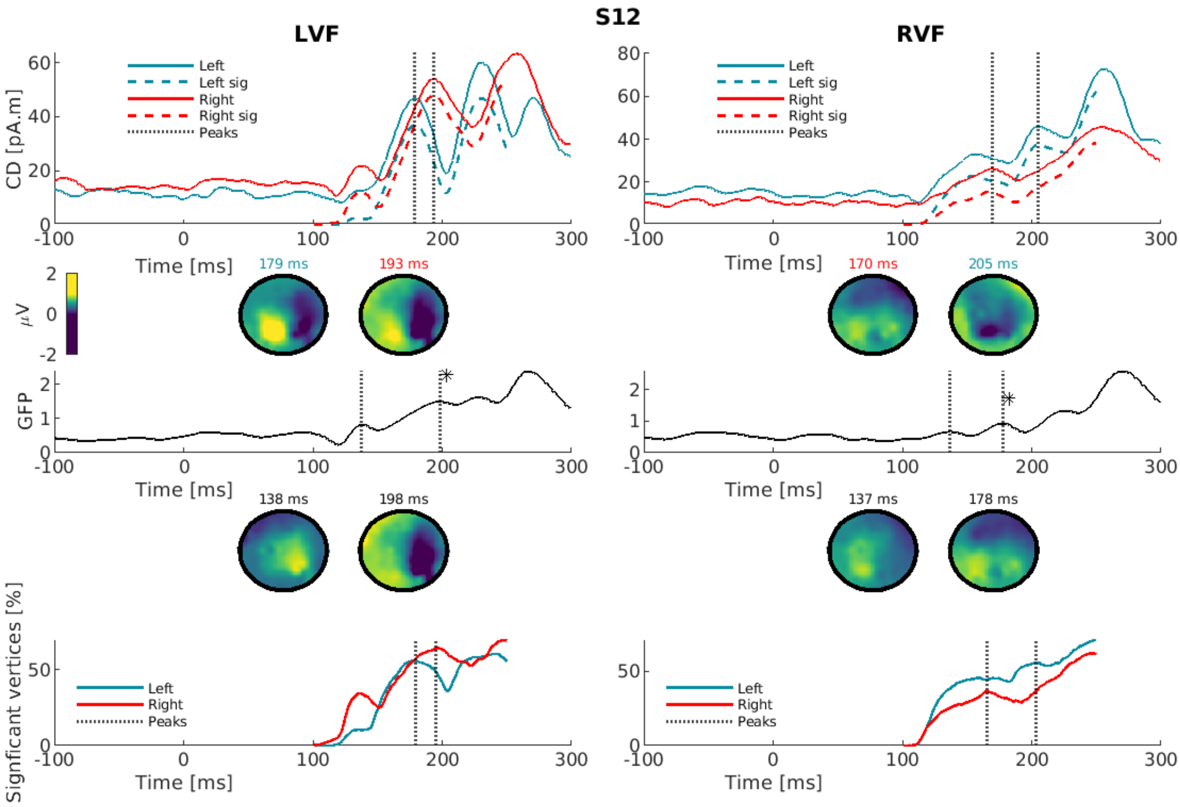
S12
LVF
RVF
CD [pA.m]
Left
Left sig
Right
Right sig
Peaks
Time [ms]
179 ms
193 ms
170 ms
205 ms
μV
GFP
138 ms
198 ms
137 ms
178 ms
Signficant vertices [%]

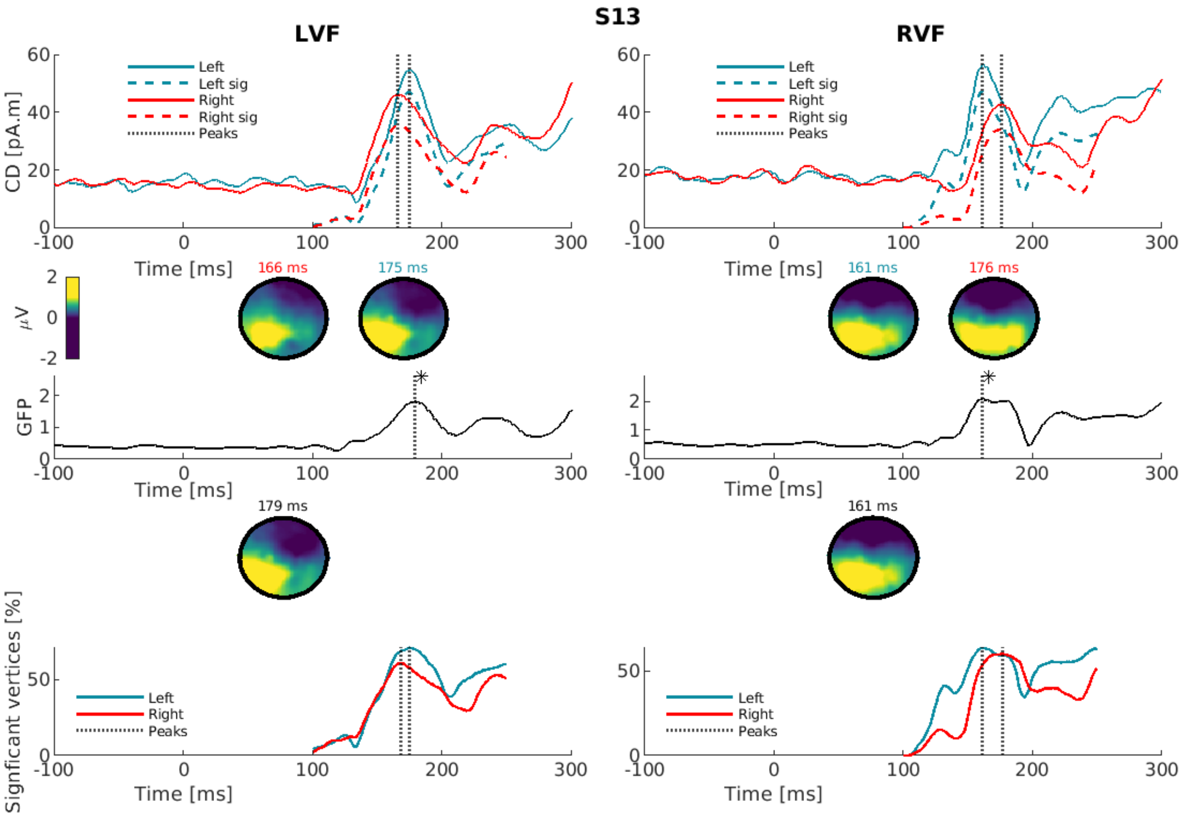
S13
LVF
RVF
Left
Left sig
Right
Right sig
Peaks
CD [pA.m]
Time [ms]
166 ms
175 ms
161 ms
176 ms
µV
GFP
179 ms
161 ms
Signficant vertices [%]

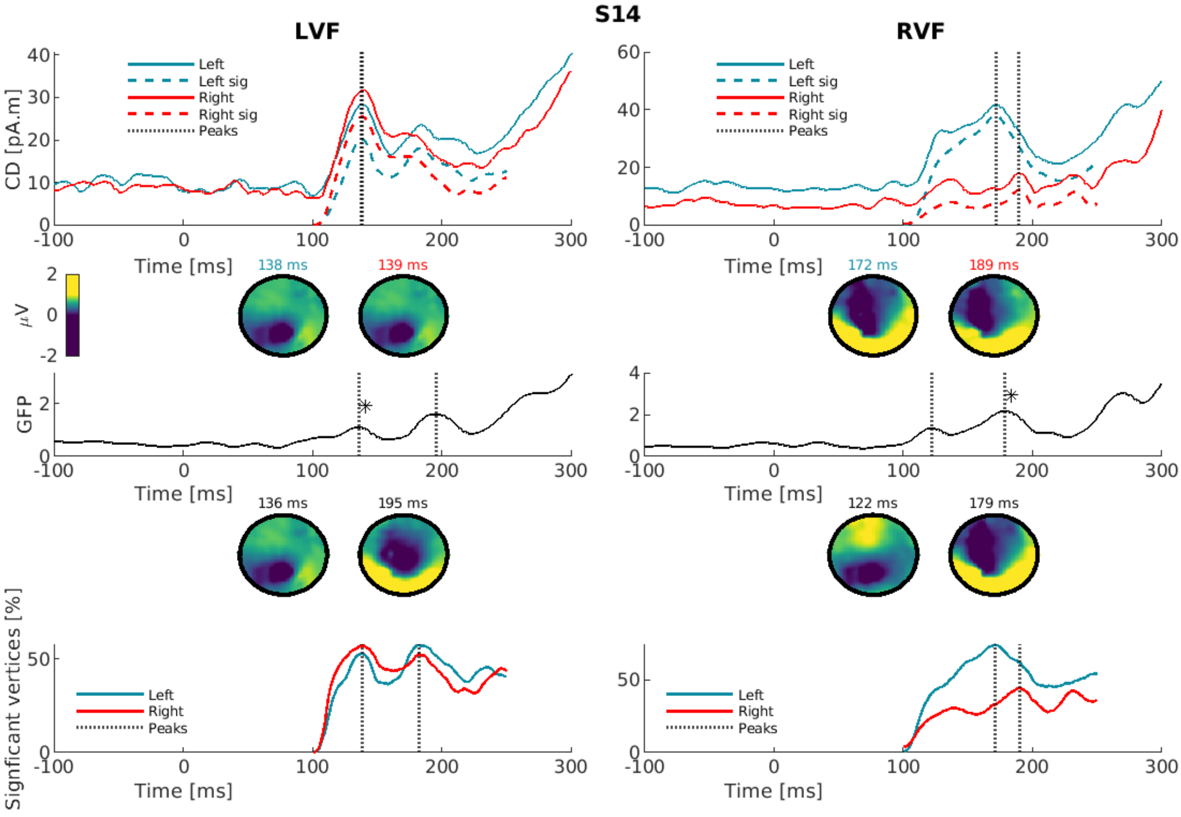
S14
LVF
RVF
Left
Left sig
Right
Right sig
Peaks
CD [pA.m]
Time [ms]
138 ms
139 ms
172 ms
189 ms
µV
GFP
136 ms
195 ms
122 ms
179 ms
Signficant vertices [%]

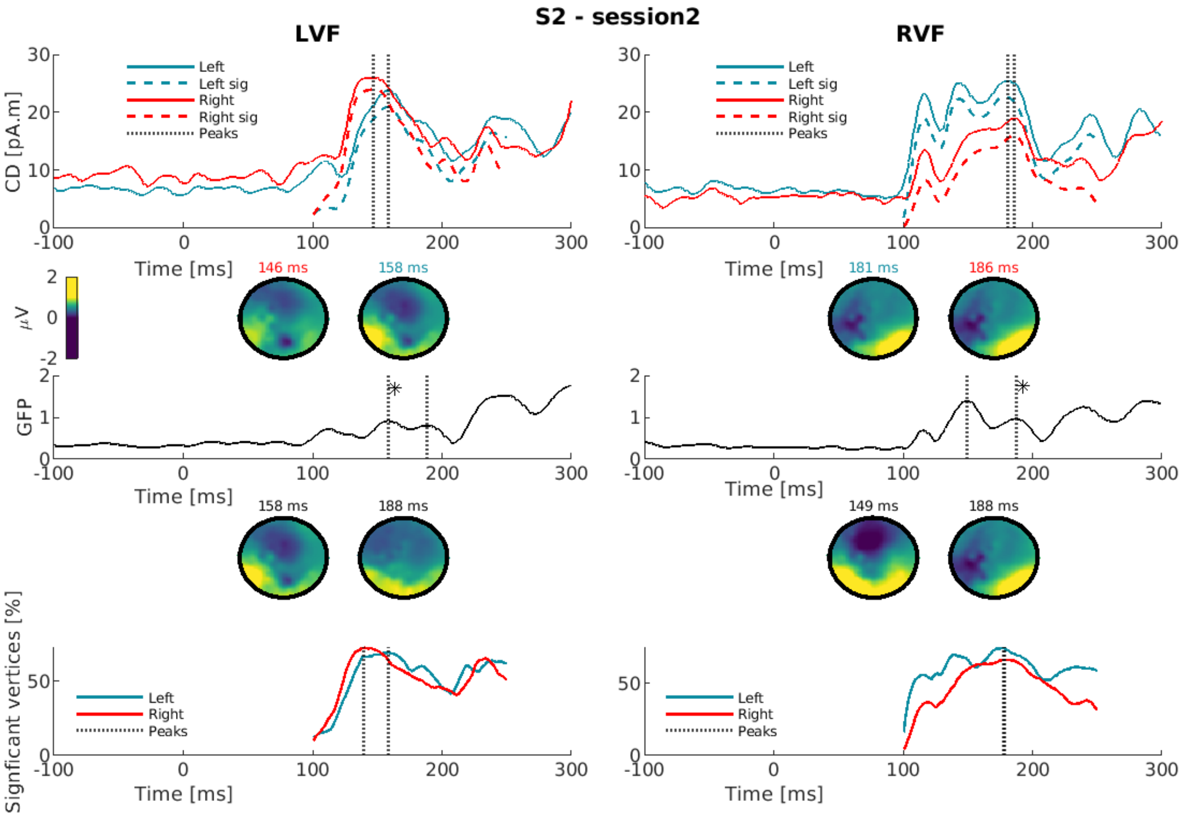
S2 - session2
LVF
RVF
Left
Left sig
Right
Right sig
Peaks
CD [pA.m]
Time [ms]
146 ms
158 ms
181 ms
186 ms
μV
GFP
158 ms
188 ms
149 ms
188 ms
Signficant vertices [%]

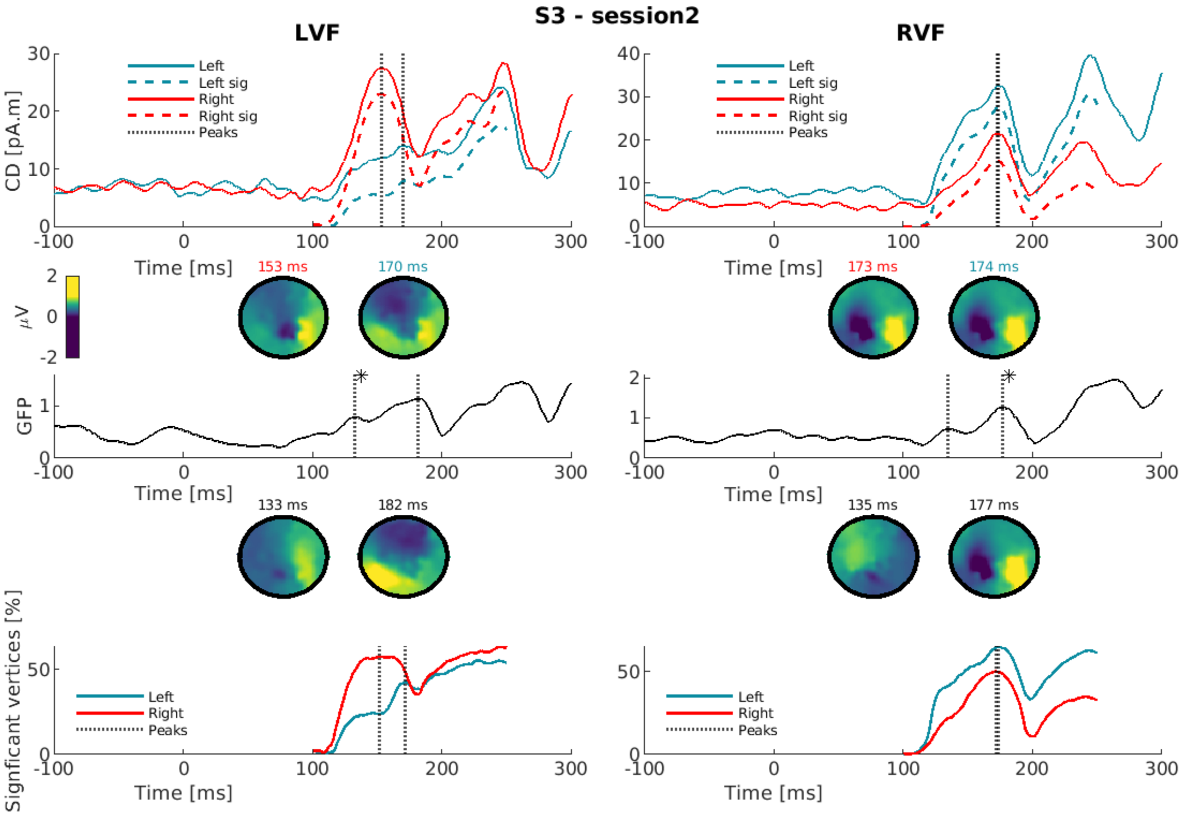
S3 - session2
LVF
RVF
Left
Left sig
Right
Right sig
Peaks
CD [pA.m]
Time [ms]
153 ms
170 ms
173 ms
174 ms
μV
GFP
133 ms
182 ms
135 ms
177 ms
Signficant vertices [%]

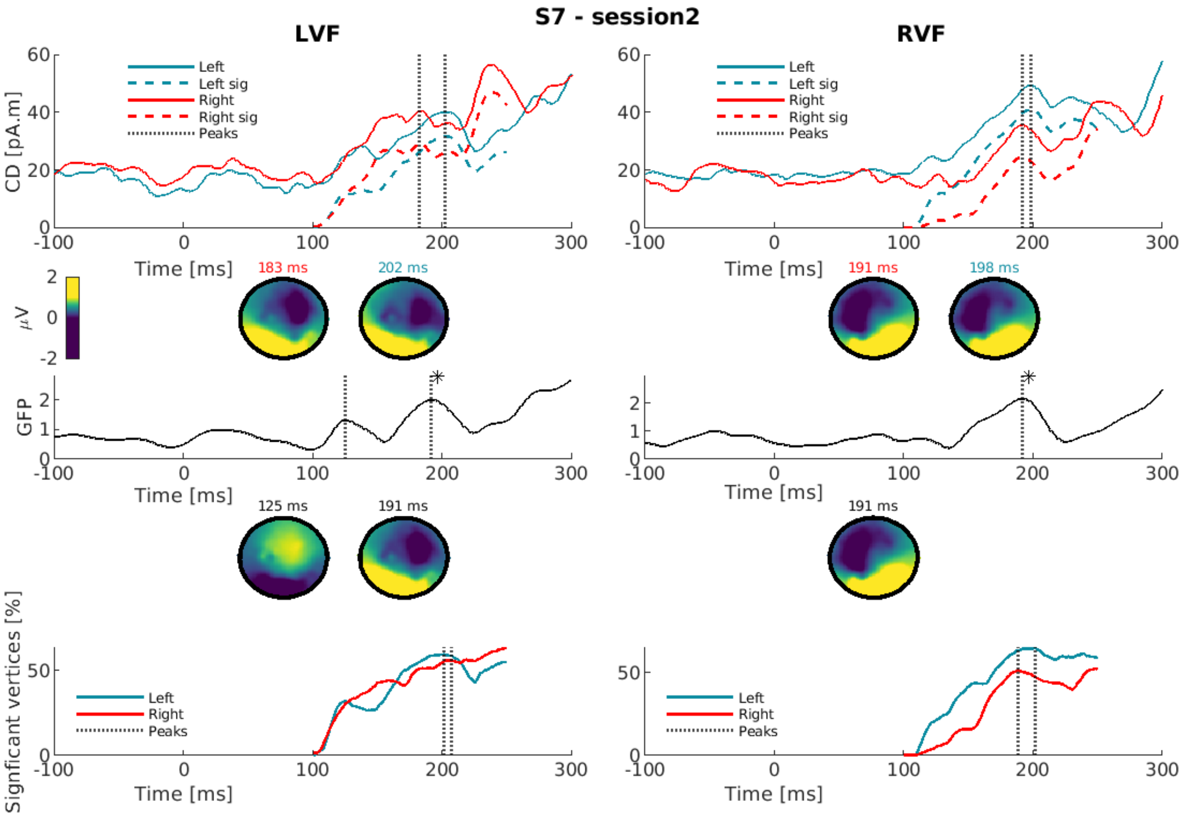

S7 - session2
LVF
RVF
CD [pA.m]
Left
Left sig
Right
Right sig
Peaks
Time [ms]
183 ms
202 ms
191 ms
198 ms
μV
GFP
125 ms
191 ms
Signficant vertices [%]

<u>Supplementary Fig. 3:</u>

Impact of inaccurate input parameters on the estimated morphological features. A 10% bias increase in conduction velocity leads to an average bias of ~22% and ~2% for θ and $\beta$, respectively, within the plausible range of in-vivo data values (0.01<θ<0.9 $\mu$m).

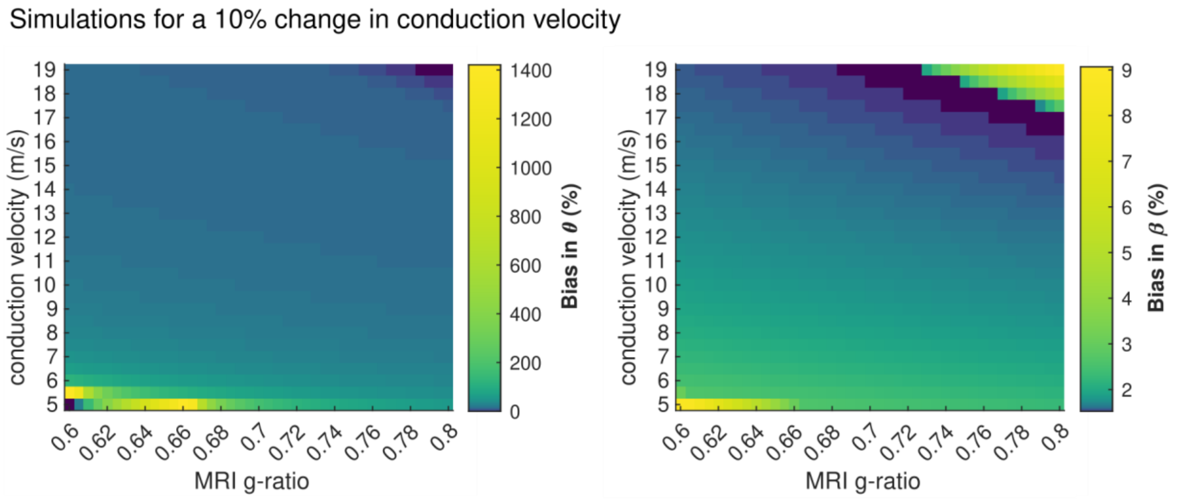


<u>Supplementary Table 1:</u>

IHTT estimates for the LVF and RVF with the three metrics for the 14 subjects obtained with the first amplitude maximum and with the second amplitude maximum if existed. Time in ms.

| | **Left Visual Field (LVF)** | | | **Right Visual Field (RVF)** | | |
|---|---|---|---|---|---|---|
| | **Original CDs** | **Masked CDs** | **Number of vertices** | **Original CDs** | **Masked CDs** | **Number of vertices** |
| **S1** | 35 | 36 | 46; 19 | 22 | 23 | 29 |
| **S2** | 12 | 11 | 9 | 42 | 42 | 43; 5 |
| **S3** | -5; 23 | -7; 24 | -18; 23 | 5 | 4 | 6 |
| **S4** | 29; -15 | 31 | 33; -5 | -1 | -1 | 7 |
| **S5** | 35 | 37; 12 | -1 | 19 | -7; 15 | -13 |
| **S6** | 2 | 4 | 9 | 56 | 57 | 13; 59 |
| **S7** | 36 | 35 | 32 | -10 | -11 | -16; -3 |
| **S8** | 47 | 50 | 51; 32 | -20 | -21 | -22; 18 |
| **S9** | 49 | 49 | 51 | -13 | -14 | -8 |
| **S10** | 15 | 13 | 13; -19 | 25; -4 | 22; -6 | 20; 2 |
| **S11** | 13 | 15 | 14 | 21 | 22 | 16; 14 |
| **S12** | -15 | -15 | -16 | -35 | -39 | -38 |
| **S13** | 9 | 7 | 7 | 15 | 15 | 16 |
| **S14** | -1 | 0 | 44 | 18 | 18 | 20 |
